# Differentiating erythroblasts adapt to turbulent flow by accelerating maturation and activating cholesterol biosynthesis

**DOI:** 10.1101/2023.12.08.570773

**Authors:** Giulia Iacono, Asena Abay, Joan S. Gallego Murillo, Francesca Aglialoro, Nurcan Yagci, Kerly Fu, Arthur F. Svendsen, Marieke von Lindern, Emile van den Akker

## Abstract

*In vitro* culture of erythroblasts (EBL) and production of mature erythrocytes for transfusions requires upscaling in fluidic-turbulent bioreactors, resulting in membrane shear stress. For the implementation of erythroid cultures in bioreactors, understanding the effects of mechanical stress on terminal EBL differentiation is required. To this end, we investigated the effect of orbital shaking-induced shear stress on differentiating CD49d^+^CD235^low^ primary human EBL towards enucleated reticulocytes at the molecular, cellular, and functional level. Orbital shaking at the onset of EBL differentiation enhanced cell maturation increasing enucleation percentage compared to static cultures, without cell viability loss. Transcriptome analysis uncovered 505 genes differentially expressed between static and dynamic cultures, with genes involved in lipid and cholesterol biosynthesis upregulated in dynamic conditions. In line with this, cells differentiated in orbital-shakers showed increased cholesterol concentration and osmotic resistance compared to static cultures. HMGCR (3-Hydroxy-3-Methylglutaryl-CoA-Reductase), rate-limiting enzyme of the cholesterol biosynthesis pathway, showed earlier and significantly higher induction during differentiation in dynamic. The severe loss of EBL in dynamic, but not in static conditions, due to HMGCR inhibition confirmed the ability of EBL to adapt to shear stress through modulating of their transcriptional program and upregulation of cholesterol biosynthesis. This work sheds light into specific mechanisms that will assist the successful upscaling of erythroid differentiation in turbulent bioreactors. In addition, as shear-stress on hematopoietic cells is also occurring within the bone marrow niche, these results introduces a potential novel signalling axis that need to be integrated into the known transduction pathways that control erythropoiesis.

## Introduction

Red blood cell (RBC) transfusion is the most common form of cell therapy, yet the practice has remained relatively unchanged for decades.^1^ Although life-saving, this donor-dependent process carries risks such as alloimmunization and spreading of blood-borne diseases. Thus, the medical need for large-scale production of erythrocytes lacking immunogenic blood group antigens is evident.^2^ Moreover, cultured RBCs (cRBC) hold great therapeutic potential as carriers for therapeutic molecules such as enzymes or inducers of tolerance.^3^

The production of functional, enucleated red blood cells is feasible in both static and turbulent environments.^4–8^ Nevertheless, the upscaling of RBC cultures to produce clinically relevant numbers in a cost-effective manner still remains a challenge.

Stirred bioreactors are effective for large scale erythroid cultures, similar to upscaling efforts for other mammalian cells.^4,9–11^ However, due to the turbulent nature of the flow in stirred bioreactor settings, cells experience a range of distinct mechanical forces.^4^ These forces will greatly vary depending on factors like suspension density, mixing speed and/or fluid volume, and air sparging strategies,^12^ and may act on mechanical sensors that are expressed and functional during erythropoiesis.^13–16^ Downstream these mechano-sensors, specific signal transduction pathways can be induced, including MAPKs, STAT5, NFATs that regulate erythroblast (EBL) specification from hematopoietic stem cells,^15^ perturbs erythroblast proliferation potential,^14^ regulates integrin activation^13^ and reticulocyte maturation^17^. Here, we investigated how shear stress affects terminal erythroblast differentiation using orbital shakers that are widely used in culture of other mammalian cells and hence have been extensively investigated in terms of fluid dynamics and biological effects of mechanical forces.^18–20^ The RBC plasma membrane consists of a lipid bilayer with embedded proteins attached to the underlying membrane skeleton. A variety of lipids and phospholipid species represent the core structure of this membrane among which cholesterol is the most abundant.^21^ Cholesterol, and particularly the ratio between cholesterol and phospholipids, controls cell membrane rigidity and fluidity.^22^ Cholesterol concentration in RBC plasma membranes affects cell morphology and functionality. Increased plasma membrane cholesterol affects RBC shape, impairs RBC flow properties, and influences plasma membrane functions such as permeability and transport.^22,23^ Moreover, cell viability and fragility in turbulent conditions is highly dependent on plasma membrane fluidity (PMF). Eukaryotic cells are capable of adaptive responses to changing membrane lipid content to ensure optimal PMF by *de novo* lipid biosynthesis.^24^ Interestingly, it has been shown that lipid membrane composition changes in shear flow environments, which has been linked to mechanosensing.^25^

Here, we show that erythroid differentiation is accelerated upon orbital shaking, without affecting cell viability. Adaptation to shear stress is reflected by an altered gene expression profile including upregulation of cholesterol metabolism pathways. We uncovered that a generally regarded rate-limiting enzyme, 3-Hydroxy-3-Methylglutaryl-CoA Reductase (HMGCR) of the cholesterol synthesis plays a central role in this phenomenon.

This data provides the first analysis of the (transcriptional) program induced by culturing erythrocyte precursors in shear stress environments. Importantly, this response was not detrimental to erythroid differentiation but changed RBC functionality and membrane properties and was paramount to the adaptive response to shear environments. Taken together, this study provides new insights that will assist in the upscaling of cRBC production, which can be employed for clinical use.

## Material and Methods

### Erythroblast Expansion and Differentiation

Peripheral blood mononuclear cells (PBMC) from healthy donors were isolated by ficoll density separation according to the manufacturer’s protocol (GE Healthcare, Chicago, Illinois, USA). Informed consent was given in accordance with the Declaration of Helsinki, the Dutch National and Sanquin Internal Ethic Boards. PBMC were cultured to proerythroblasts in IMDM medium (PAN P04-20250) supplemented with EPO (2U/mL; ProSpec, East Brunswick, NJ, USA), human recombinant Stem Cell Factor (100ng/mL, supernatant SCF producing cell line) and dexamethasone (1μM; Sigma, St. Louis, MO, USA).^5^ EBL cultures were kept at a density of 0.7-1.5 million/mL by dilution with fresh medium. To differentiate EBL to reticulocytes, cells were washed twice in PBS, to remove any residual growth factors, and resuspended in differentiation media (1 million/mL): IMDM supplemented with EPO (10U/mL), 5% Omniplasma (Octopharma, Wien, Austria), holotransferrin (700µg/mL; Sanquin), and heparin (5U/mL; LEO Pharma BV, Breda, The Netherlands). Cell counts were obtained using a CASY (CASY®Model TTC, Schärfe System GmbH, Reutlingen, Germany). The day of change to differentiation media is referred to as Day 0 of differentiation.

### Orbital shaking culture conditions

Orbital shaking was performed using 125mL Corning Erlenmeyer cell culture flasks with ventilated caps (Sigma Aldrich, Munich, Germany) on a Thermo Scientific™ CO2 Resistant Shaker (Catalog N°: 15341105). For all experiments 250 revolutions per minute (RPM) was used, yielding to a 1.4 Pa maximum wall shear stress. The starting volume of dishes and flasks samples was of 12mL and 25mL respectively. Conditions were switched as indicated.

### Flow Cytometry

Samples were resuspended in HEPES buffer (132mM NaCl, 20mM HEPES, 6mM KCl, 1mM MgSO4, 1.2mMK2HPO4) supplemented with 0.5% Bovine Serum Albumin (BSA) and incubated with the following antibodies for 30 minutes: antiCD235a-PE (OriGene Technologies, Rockville, MD, USA), antiCD49d-PB (BD Biosciences, San Jose, CA, US), TO (Thiazole orange-Sigma), antiCD71-PB (Miltenyi Biotec), PI-PE (Propidium Iodide - Invitrogen) and DRAQ5 (1ng/mL, incubation 5 minutes at RT; BioLegend, San Diego, CA, USA). Samples were measured using a FACS Canto II (BD Biosciences, Oxford, UK) and results were analysed using FlowJo software (FlowJo v10; Tree Star, Inc., Ashland, OR USA).

### Cytospin Images

Cytospins were performed with Shandon Cytospin II (Thermo Scientific), stained with benzidine (staining heme) and Differential Quik Stain Kit (PolySciences, Warrington, PA) to differentiate globin and DNA as described before^5^. Stained cytospins were imaged with 50x objective on Leica DM-2500 microscope (Leica, Germany).

### RNA-Sequencing and Analysis

EBL differentiated in static (dish) or dynamic conditions (flask) were sampled at day 1, 2 and 4 following differentiation induction. RNA was isolated from cell pellets using Trizol as indicated by the manufacturer’s protocol (Invitrogen Technologies, CA, USA). DNA remnants were removed by RNA clean up kits (Zymo Research, CA, USA). RT-reactions and RNA sequencing was performed by GENEWIZ (NJ, USA) using random hexamers, with on average 30 million reads deep. After sequencing, quality control and adapter trimming were performed using FastQC v0.11.9 (Babaham Bioinformatics) and fastp 0.23.2^26^ with additional arguments -- length_required 36, --cut_right, --cut_right_window_size 4, --cut_right_mean_quality 15. Next, pair-ended trimmed fastq files were mapped to GRCh38.p13 (Ensembl 87) using STAR 2.7.0f^27^ with --quantMode GeneCounts. Deferential expression analysis was performed using second stranded reads in EdgeR 3.40.2.^28^ In short, low expressed reads were filtered out, all samples were TMM normalised, and dispersion was calculated using the estimateDisp function. Samples were categorised in two different groups (dish and flask) for the different differentiation days. Pairwise comparisons between the different groups were carried out for the different days using the glmLRT/ glmLRT functions; p-values were bonferroni adjusted and genes with adjpval ≤ 0.05 were considered differentially expressed between groups. Analyses were performed in Ubuntu 22.04.2 LTS and R version 4.2.3. Heatmaps were plotted using Morpheus (https://software.broadinstitute.org/morpheus). Gene ontology analysis was performed using EnRicHR.^29–31^

### Western blotting

EBL were differentiated as indicated above. Cells were sampled daily, pelleted and washed 3 times with ice-cold PBS followed by lysis in CARIN lysis buffer (20mM Tris-HClpH 8.0, 138mM NaCl, 10mM EDTA, 100mM NaF, 1% Nonidet P-40, 10% glycerol). Lysates were boiled in Laemmli sample buffer (2% sodium dodecyl sulfate [SDS] wt/vol, 10% glycerol, 1% 2- mercaptoethanol, 60mM Tris-HCl pH6.8, brome-phenol blue) for 5 minutes at 95°C, subjected to SDS-polyacrylamide gel electrophoresis, blotted using iBlot-PVDF blotting system (ThermoFisher Scientific, Bleiswijk, The Netherlands), and stained as indicated in the figure legends: anti-HMGCR (ab242315, Abcam) and anti-GAPDH (MAB374, Millipore).

### Stability and deformability assays

On day 12 of differentiation cell obtained from static (dish) and dynamic cultures (flask) were filtered using Spuitfilters (Acrodisc® WBC #514-0682) to remove nucleated cells and pyrenocytes. Purified reticulocytes were washed 3 times with PBS. To measure the membrane’s stability, reticulocytes were subjected to osmotic shock by incubation in 0%, 0.5%, 0.6% and 0.7% NaCl for 5 minutes at RT. Cells were centrifuged, the supernatant’s absorbance measured at 415nm. Reticulocytes’ deformability was assessed using ARCA (ARCA-Linkam CSS450)^32^: cells were suspended in PVP buffer (5% Polyvinylpyrrolidone in PBS) and exposed to a shear stress of 3 Pa and 10 Pa.^5^

### Cholesterol quantification

2×10^6^ cells differentiated in static (dish), or dynamic (flask) conditions were washed 3 times with PBS and lysed in chloroform:isopropanol:NP40 (7:11:0.1). Lysates were centrifuged and the organic phase air dried. The leftover volume was subjected to a speed vacuum for 45 minutes. Total cholesterol amount was measured using Cholesterol Quantitation Kit (Sigma, mak043-1kt) and the sample’s absorbance measured at 570 nm.

### HMGCR inhibition

On day 0 of differentiation, EBL were incubated with different concentrations of the HMGCR inhibitor Lovastatin (ab120614, Abcam: 0.1µM, 0.3µM, and 1µM). Cells were differentiated in static (dish) and dynamic (flask) conditions. Cells differentiation, enucleation, stability and deformability were assessed at the end of differentiation as described above.

### Reticulocytes enrichment

Reticulocytes were isolated starting from 3 ml of packed peripheral blood RBC using CD71-MACS beads (Miltenyi 130-046-201) according to the manufacturer protocols. Reticulocyte enrichment was evaluated by TO (Thiazole orange) and anti-CD71 flow cytometry staining.

## Data Sharing Statement

For original data, please contact.

Sequencing data are available at GEO under accession number GSE247121

Upon request, the raw data supporting the conclusion of this article will be made available by the authors without undue reservation.

## Results

### Orbital Shaking Accelerates Erythroid Maturation

In static conditions, EBL differentiation to enucleated reticulocytes is accompanied by a progressive loss of CD49d and an increase in CD235a expression, taking approximately 10 days (Figure 1A). We exposed EBL to orbital shaking to generate an average shear stress of 1.4 Pa, comparable to the shear stress at the point of the impeller of the bioreactors used^4^ (estimation based on Odeleye^33^). Both static and dynamic cultures contained similar CD49d^-^ CD235a^+^ population at the end of the differentiation, however dynamic conditions showed significantly faster differentiation kinetics, reaching ∼90% CD49d^-^/CD235a^+^ cells 4 days earlier compared to static cultures (Figure 1A, B). A significant increase in enucleated reticulocytes (DRAQ5^-^ cells) was observed at day 4 and 6 in dynamic cultures compared to static cultures, which eventually progressed to similar enucleation percentages at the end of culture (Figure 1C, D; Figure S1A). Faster differentiation was further confirmed by morphological analysis (Figure S1B). At the end of differentiation, dynamic cultures demonstrated a slight decrease – yet not significant – in EBL and reticulocytes production (Figure 1E). Together these results indicate that orbital shaking leads to accelerated differentiation of EBL to enucleated reticulocytes.

**Figure 1.**
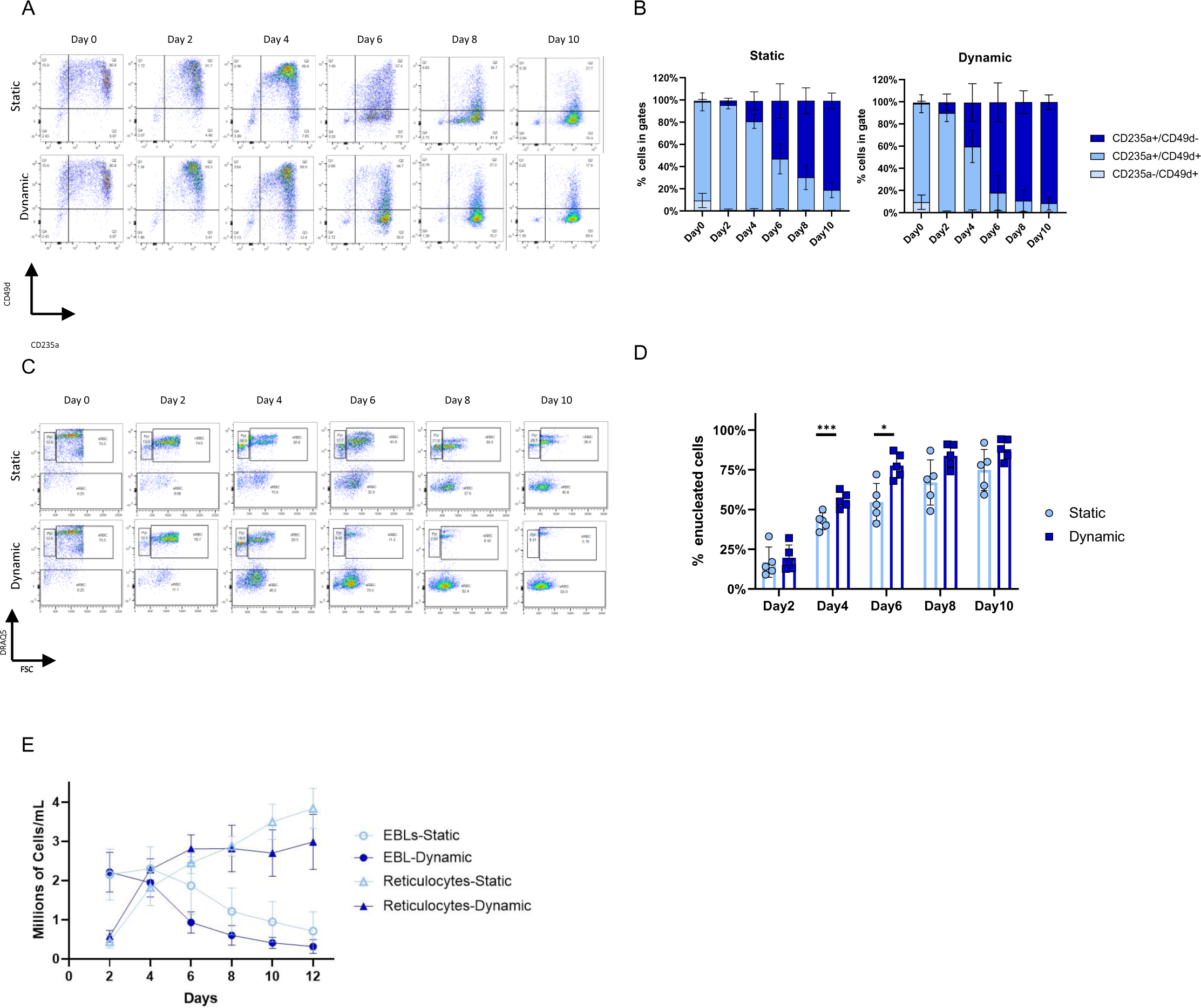
Orbital shaker accelerates maturation and enucleation compared to static cultures. Erythroblasts (EBL) were differentiated for up to 10 days in dishes (static) or orbital shakers (dynamic). A) Erythroid cells were harvested at indicated days from dishes and flasks and stained with anti-CD49d (Alexa Fluor 405, y-axis) and anti-CD235a (PE, x-axis), and analysed by flow cytometry. B) Maturation states in static vs. dynamic conditions as explained in (A) were averaged for 5 donors (n=5) by quantifying the percentage of cells within the respective gates Q1: CD49d^+^/CD235^-^ immature EBL, Q2: CD49^+^/CD235a^+^ EBL, and Q3: CD49^-^/CD235a^+^ late EBL and reticulocytes. C) Enucleation was quantified by staining cells with the cell-permeable DNA-dye DRAQ5 (APC, y-axis), forward scatter (FSC, x-axis) on the indicated days. Representative scatter plot displaying the gating for reticulocytes, EBL, and pyrenocytes is shown in supplemental figure 1A. Enucleation percentage is defined as the %reticulocyte/ (%reticulocytes + %nucleated cells) using the gates as defined in supplemental figure 1A. D) Enucleation states in static vs. dynamic conditions as explained in (C) were averaged for 5 donors (n=5) by quantifying the percentage of cells within the respective gates: reticulocytes, erythroblasts, and pyrenocytes for 5 donors (n=5). E) Cell-counts for reticulocytes and EBL that were cultured in static or dynamic conditions for 4 donors (n=4). The distinction between the two populations was done via DRAQ5 staining. B, D and E) Paired two-tailed student t-test was performed, with ** p<0.01 and *p<0.05. Unless marked, no significance was observed.

### Orbital shaking accelerates erythroblast differentiation immediately after differentiation induction

Next, we investigated whether dynamic conditions expedite differentiation at a specific EBL maturation stage. Differentiating EBL were switched between static and dynamic conditions on different time points and the enucleation rate of the cultures was measured. In line with Figure 1, an inverse correlation between enucleated cells and the initial dynamic culture time was observed (Figure 2).

**Figure 2.**
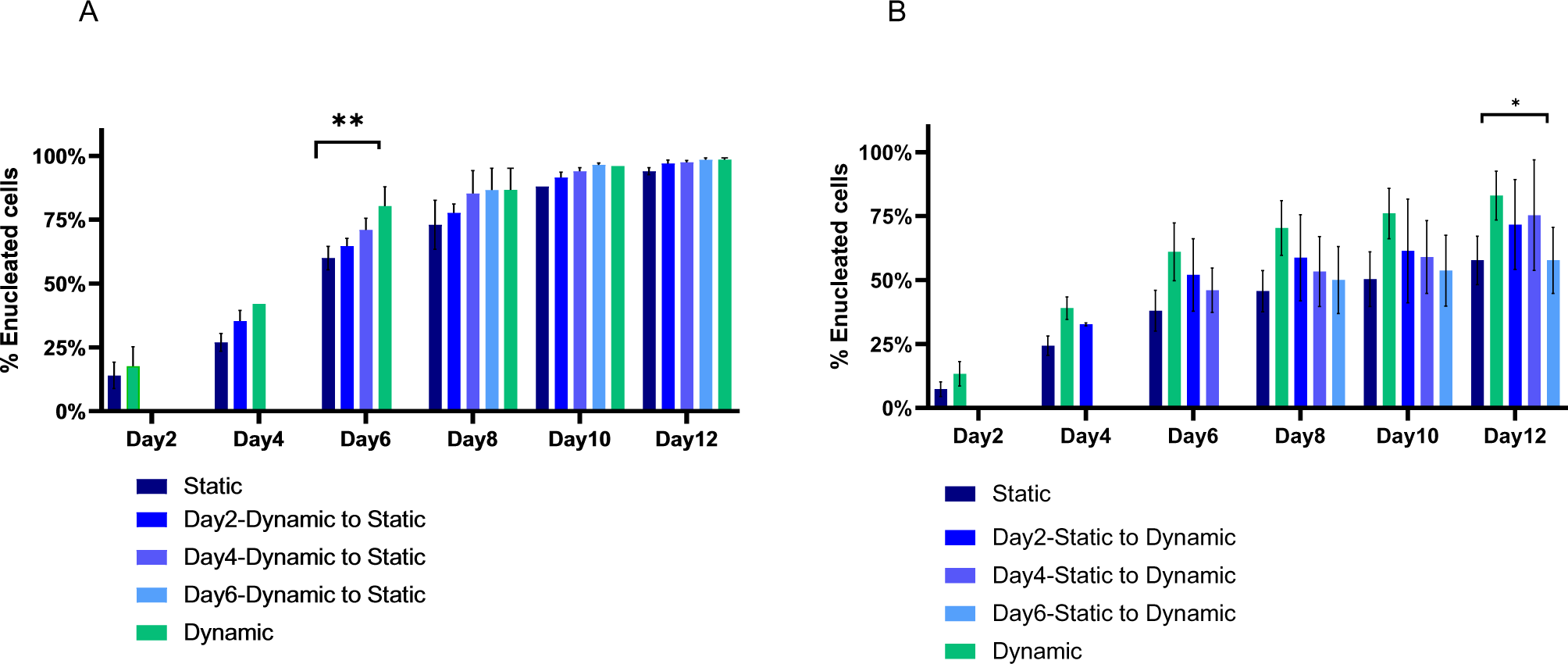
Shear-induced enhanced maturation is progressively lost during erythroblast differentiation. A) Enucleation percentage (calculated as described for figure 1C) of samples switched from dynamic to static culture on Day 2, 4, or 6 of differentiation, or not switched, for 3 donors (n=3). B) Enucleation percentage of samples switched from static culture to dynamic culture on Day 2, 4 or 6 of differentiation, as well as dish (static) and flask (dynamic) controls averaged for 3 donors (n=3). Paired two-tailed student t-test was performed, with ** p<0.01 and *p<0.05. Unless marked, no significance was observed.

Cultures switched from orbital shaking to static on day 2 showed slower enucleation kinetic compared to the samples switched on day 4. Importantly, all conditions reached comparable levels of enucleation indicating that the static cultures are not blocked in differentiation but that orbital shaking accelerates the process (Figure 2A).

In the reverse experiment, transferring static cultures to dynamic conditions on day 2 and day 4 accelerated enucleation compared to the static control, but still lagged behind compared to cells under continuous orbital shaking. Switching at day 6 to orbital shaking conditions did not accelerate enucleation suggesting that the regulation of accelerated differentiation occurs before day 6 (Figure 2B). Together the data suggest that accelerated differentiation upon orbital shaking is regulated during the early phase of EBL differentiation between day 0 and 4.

### Shear stress alters the erythroid transcriptome

To dissect the mechanism whereby dynamic conditions show accelerated differentiation, we performed transcriptomic analysis of both conditions for the first 4 days of culture.

Principal Component Analysis (PCA) demonstrated samples cluster per differentiation condition and time, where day 4 demonstrated the biggest difference (Figure S2A). Differential expression analysis between groups demonstrated many differentially expressed genes (DEGs) upon EBL differentiation, with 505 DEGs at day 4 of differentiation. (Figure S2B, C, Table S1-3).

Pearson hierarchical and K-means clustering of these DEGs revealed 3 different gene expression profiles, assigned to 3 major clusters (Figure 3A). Genes assigned to the first cluster (K1), showed an overall down-regulation during differentiation in both static and dynamic cultures – but the transcriptional levels were increased in dynamic conditions (Figure 3A). The second cluster (K2) is comprised of genes which are up-regulated in dynamic cultures; the third cluster (K3) represents up-regulated genes in static conditions.

**Figure 3.**
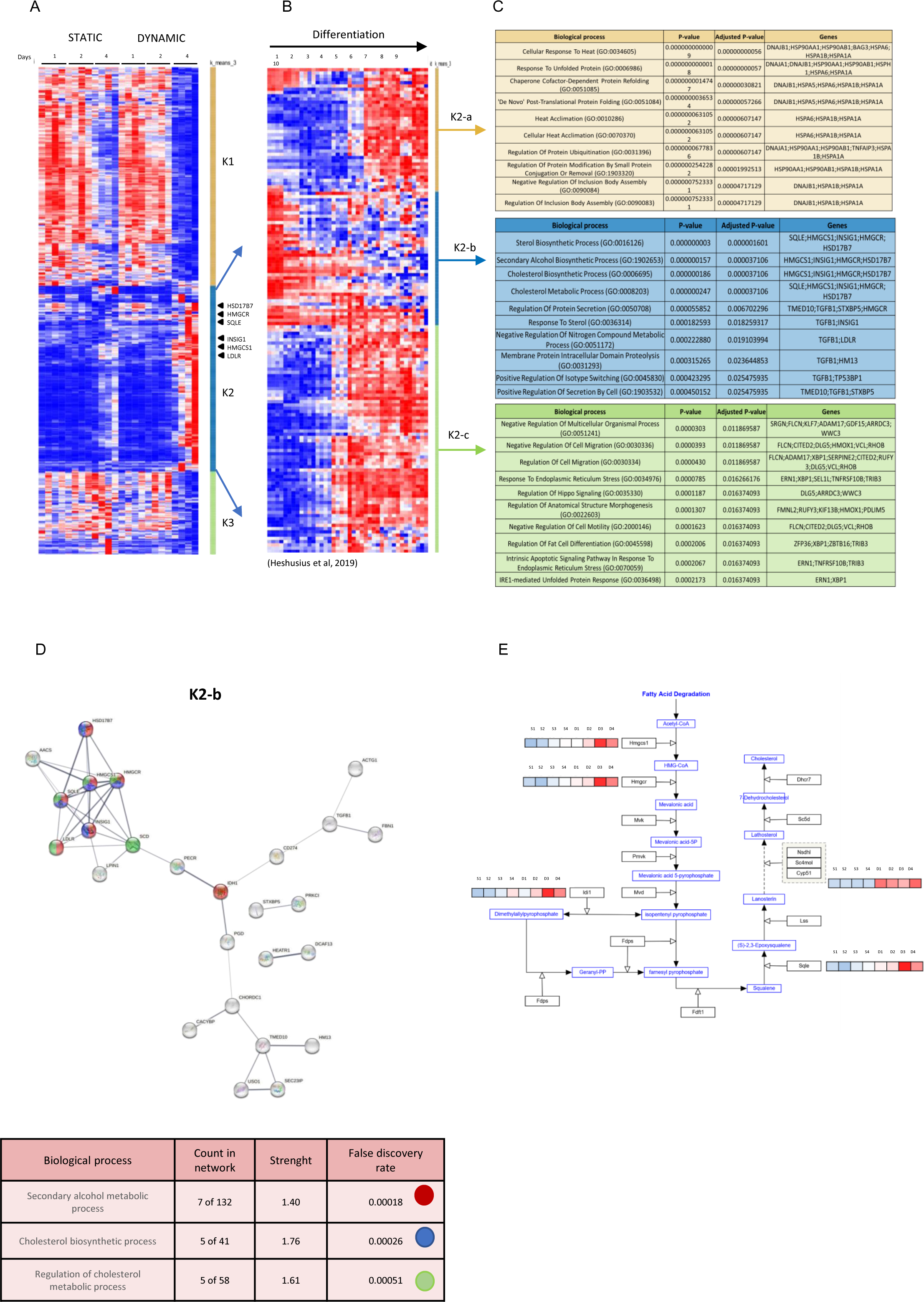
RNA analysis between static and dynamic erythroblast differentiation confirms accelerated maturation and characterises the involved processes. Erythroblasts (EBL) of four different donors (n=4) were differentiated for 4 days as indicated in material and methods. A) Heatmap showing a pearson hierarchical clustering of z-scores of the 505 differentially expressed RNAs between Day1 and Day4. K-means clustering, reveals 3 clusters indicated as K1, K2, K3. Genes involved in the cholesterol biosynthesis pathway are reported next to their specific coordinates on the heatmap. B) The gene identifiers of the upregulated RNAs of K2 from (A) were extracted and their expression dynamics through complete differentiation of EBL to enucleated reticulocytes, as previously published by Heshusius^5^, was datamined and determined. Pearson hierarchical clustering was performed on z-scores over time in days as indicated (x-axis). Kmeans clustering (K=3) was additionally performed and indicated as K2-a, K2-b and K2-c. C) ENRICHR analysis of K2-a to K2-c from (B) shows the top 10 enriched Gene Ontology (GO) term biological processes that are associated with the RNAs in the indicated Kmeans cluster according to the adjusted p-value. D) STRING analysis of the genes within K2-b was performed and genes within the first 3 biological processes according to the false discovery rate highlighted in red, blue and green. Lines thickness indicates the strength of data support. E) Representation of cholesterol biosynthesis pathway with related z-score of genes up-regulated on day4 in dynamic (D) compared to static (S) conditions.

Gene ontology analysis revealed that genes in K1 and K3 are enriched for DNA replication and nucleosome organisation processes, respectively (Figure S3A). Moreover, K3 genes during differentiation revealed a cluster of genes upregulated during differentiation (Figure S3C). The differentiation-associated gene expression profile in K1 and K3 is in line with previously reported dataset showing similar gene expression dynamics during differentiation to enucleated reticulocytes (Figure S3B, C^5^).

We then focused on K2: in contrast to K1 and K3, transcripts within K2 were down-regulated on day 4 in static conditions but up-regulated in dynamic environments. This tendency is more evident comparing transcripts of the same donor in static and dynamic conditions. To assess the expression of these genes throughout the complete differentiation to reticulocytes, we compared our data more closely to those from Heshusius^5^ and re-clustered them based on their gene expression profile, giving rise to 3 additional clusters (K2-a/b/c, Figure 3B).

The top 10 regulated biological processes according to the adjusted p-value (EnRicHR) revealed that genes in K2-a regulate response to heat and to unfolded protein and chaperone cofactor-dependent protein refolding, while genes within K2-c are involved in the regulation of cell migration and response to ER stress. In contrast, K2-b demonstrated a strong enrichment for genes mostly involved in cholesterol/steroid metabolism being up-regulated in dynamic conditions (Figure 3C). K2-b STRING analysis confirmed the presence of a cluster of genes involved in secondary alcohol metabolic and cholesterol biosynthetic processes, among which also HMGCR, a rate limiting enzyme of the cholesterol biosynthesis pathway (Figure 3D).

In all, these data confirm accelerated differentiation of EBL upon dynamic culture conditions and provide a transcriptional and possible functional footprint of downstream effects of shear-induced transcriptional program (SITP).

### Reticulocytes generated in shear stress environment show higher cholesterol concentration and cell stability

The identification of a shear stress induced transcriptional program, can potentially affect erythrocyte homeostasis. We investigated whether shear stress-induced upregulation of cholesterol metabolism pathways affects cholesterol content resulting in cell stability and deformability differences between conditions (Figure 4).

**Figure 4.**
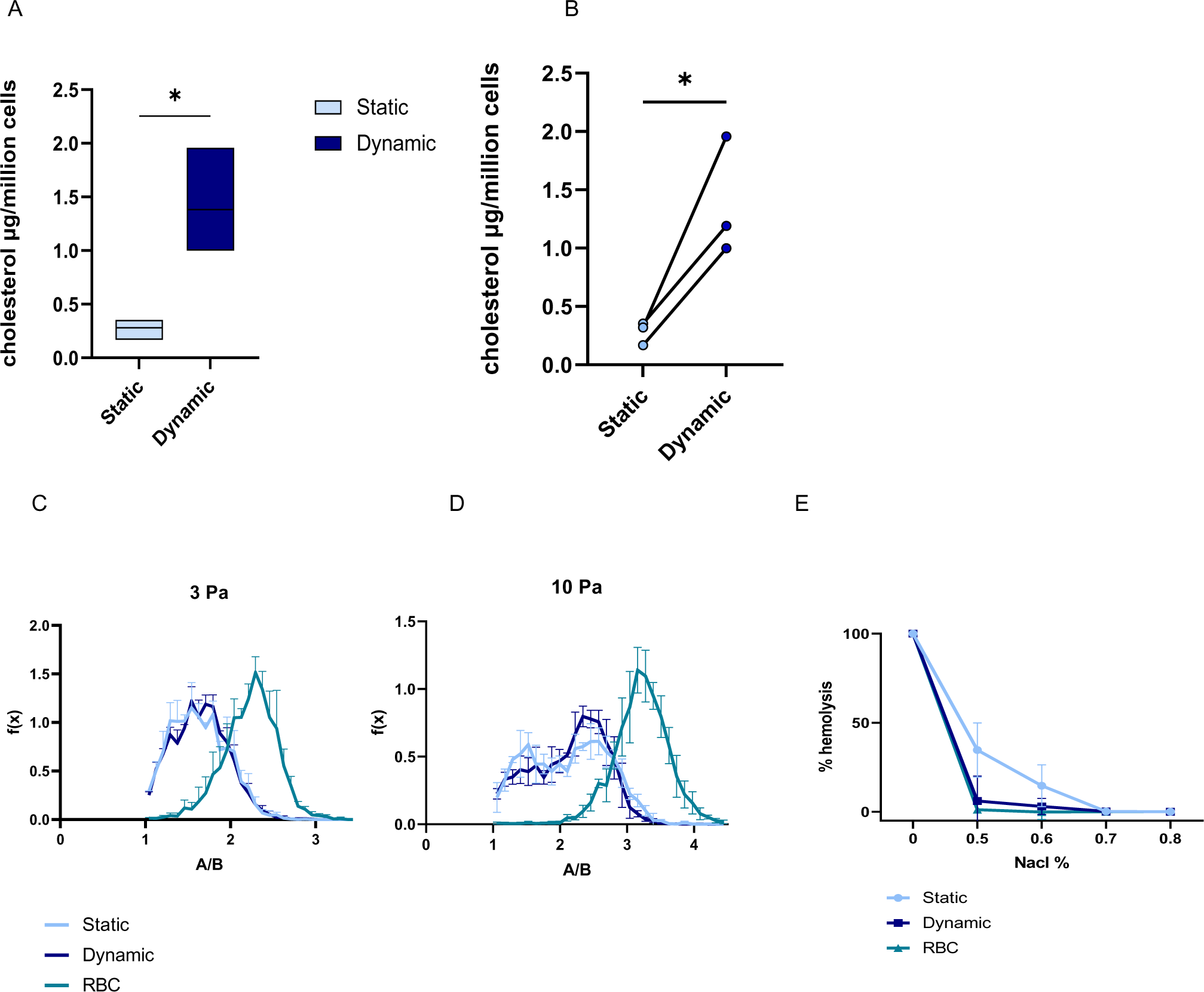
Reticulocytes obtained from dynamic cultures show higher cholesterol concentration and cell stability and similar deformability compared to reticulocytes obtained from static cultures. Erythroblasts (EBL) of three different donors (n=3) were differentiated for 12 days as indicated in material and methods. On day 12 of differentiation, part of the cells was pelleted and their cholesterol concentration was measured. Part of the cells were filtered, and the obtained reticulocytes subjected to cell deformability and stability assays. A) Cholesterol concentration in cells differentiated in static and dynamic conditions is reported in µg of cholesterol/million cells. B) Comparison of cholesterol concentration assessed as described in (A) between cells of the same donor differentiated in static vs dynamic conditions. C and D) Measure of the reticulocyte’s deformability was detected using the ARCA. Cells were subjected to a shear stress of 3 Pa (C) and 10 Pa (D). The x-axis represents the ratio between the length and the width of the cell (A/B), the y-axis is the normalised number of cells. A/B ratio directly correlates with reticulocyte deformability. E) Hemolysis percentage (y-axis) measured after incubation with different concentrations of NaCl solution (x-axis) was assessed to determine the stability of reticulocytes obtained from static and dynamic cultures. C, D and E) Native RBCs were used as control in deformability and stability assays. A and B) Paired two-tailed student t-test was performed, with ** p<0.01 and *p<0.05. Unless marked, no significance was observed.

Because the cells obtained from our cell cultures are not mature RBC but reticulocytes, we quantified cholesterol concentration in native RBC and native reticulocytes isolated from peripheral blood (Figure S4). The detected cholesterol concentration was on average 0.23 ± 0.02 µg/million and 0.13 ± 0.10 µg/million cells in RBC and reticulocytes, respectively, suggesting that the use of RBC or reticulocytes as control is interchangeable for the scope of our experiments. Cholesterol content of cells obtained from static conditions on day 12 of differentiation was on average 0.3 ± 0.09 µg/million cells, double the amount of what was previously published^34^ but comparable to the value measured in freshly isolated RBC or reticulocytes.

Corroborating with our hypothesis, cholesterol content is significantly increased in cells differentiated in dynamic conditions, reaching an average of 1.4 ± 0.5 µg/million cells (Figure 4A, B). Cholesterol controls the fluidity of cell membranes. Therefore, we measured the deformability of the cultured and freshly isolated reticulocytes using an automated rheoscope and cell analyzer (ARCA-Linkam CSS450) subjecting the cells to a shear stress of 3 Pa and 10 Pa. We detected lower deformability in reticulocytes obtained from static and dynamic cultures, compared to native RBC, but no difference was detected between static and dynamic conditions. (Figure 4C, D). In contrast, the osmotic resistance of reticulocytes derived from dynamic conditions was comparable to RBC controls where static cultures showed lower osmotic resistance (Figure 4E).

Together these data suggest that the upregulation of the cholesterol biosynthesis pathways detected in EBL differentiated in dynamic conditions is associated with higher cholesterol concentration and osmotic resistance, but similar deformability compared to cells differentiated in static conditions.

### Shear stress induces the activation of the HMGCR pathway regulating the cholesterol biosynthesis

Within the cholesterol biosynthesis pathway, the enzymes HMGCRS1 and HMGCR, IDI1, SQLE and HSD17B7 showed a similar trend to be upregulated in dynamic conditions (Figure 3D, E). Of note, HMGCR is one of the rate-limiting and fundamental coordinators of mevalonate metabolism that assures constant *de novo* synthesised cholesterol and non-sterol products.^35^

In static conditions, HMGCR expression peaked at days 4 and 5 of differentiation, where peak expression during dynamic conditions occurs during day 1 and 2 and decreased on day 3 and 4 (Figure 5A, B and Figure S5), providing further proof of the accelerated maturation in dynamic cultures. Importantly, the high expression of HMGCR detected on day 1 and day 2 of differentiation in dynamic conditions was never reached in static cultures, suggesting also an adaptation response of the cells to the shear stress and confirming the increase in gene expression observed by RNA-sequencing (Figure 4B). These data together suggest a possible connection between the upregulation of HMGCR and the increased cholesterol concentration and stability in cells differentiated in dynamic conditions.

**Figure 5.**
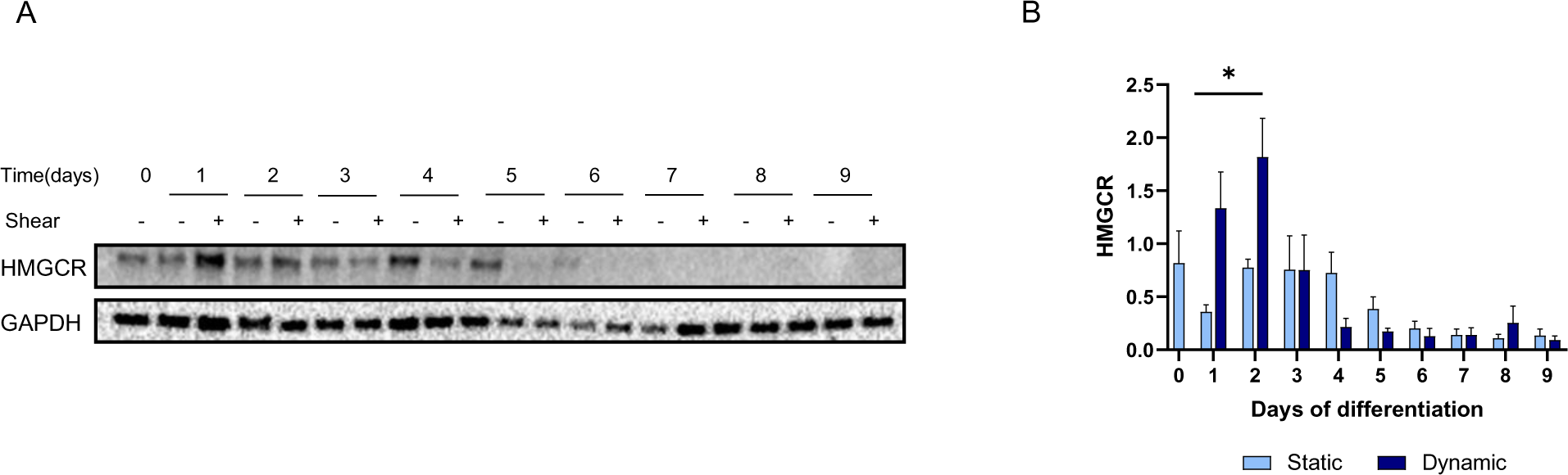
Erythroblast differentiation in shear stress environment upregulates HMGCR expression. A) HMGCR expression was assessed through western blot analysis, during erythroblasts (EBL) differentiation of 3 different donors (n=3) in static and dynamic conditions. GAPDH was used as loading control. B) Quantified expression of HMGCR during cell differentiation in dynamic and static conditions. HMGCR expression was normalised according to GAPDH expression. B) Paired two-tailed student t-test was performed, with ** p<0.01 and *p<0.05. Unless marked, no significance was observed.

### The HMGCR inhibitor lovastatin leads to a failure in adaptation to orbital shaking conditions and a severe reduction of reticulocyte formation during erythroblast differentiation

Increased production of cholesterol during dynamic conditions poses the question whether the increased HMGCR expression is essential for the adaptive response of differentiating EBL during dynamic conditions. Statins are well-described HMGCR inhibitors, used in the clinic to reduce cholesterol levels.^36^ It has been established that statins reduce RBC membrane cholesterol and increase membrane deformability and ATP release in hypercholesterolemic individuals.^37^

Lovastatin treatment at different concentrations did not affect cell yield and general hemoglobinization during EBL differentiation in static cultures. In contrast, dynamic cultures treated with the same Lovastatin concentrations showed a significant reduction of cell viability and hemoglobinization (Figure 6A, B, E and Figure S6). CD49d/CD235a marker expression was not affected in lovastatin treated EBL differentiated in static conditions, except for a reduction of CD235a^+^/CD49d^-^ cells with high (1µM) lovastatin concentrations (Figure 6C, D).

**Figure 6.**
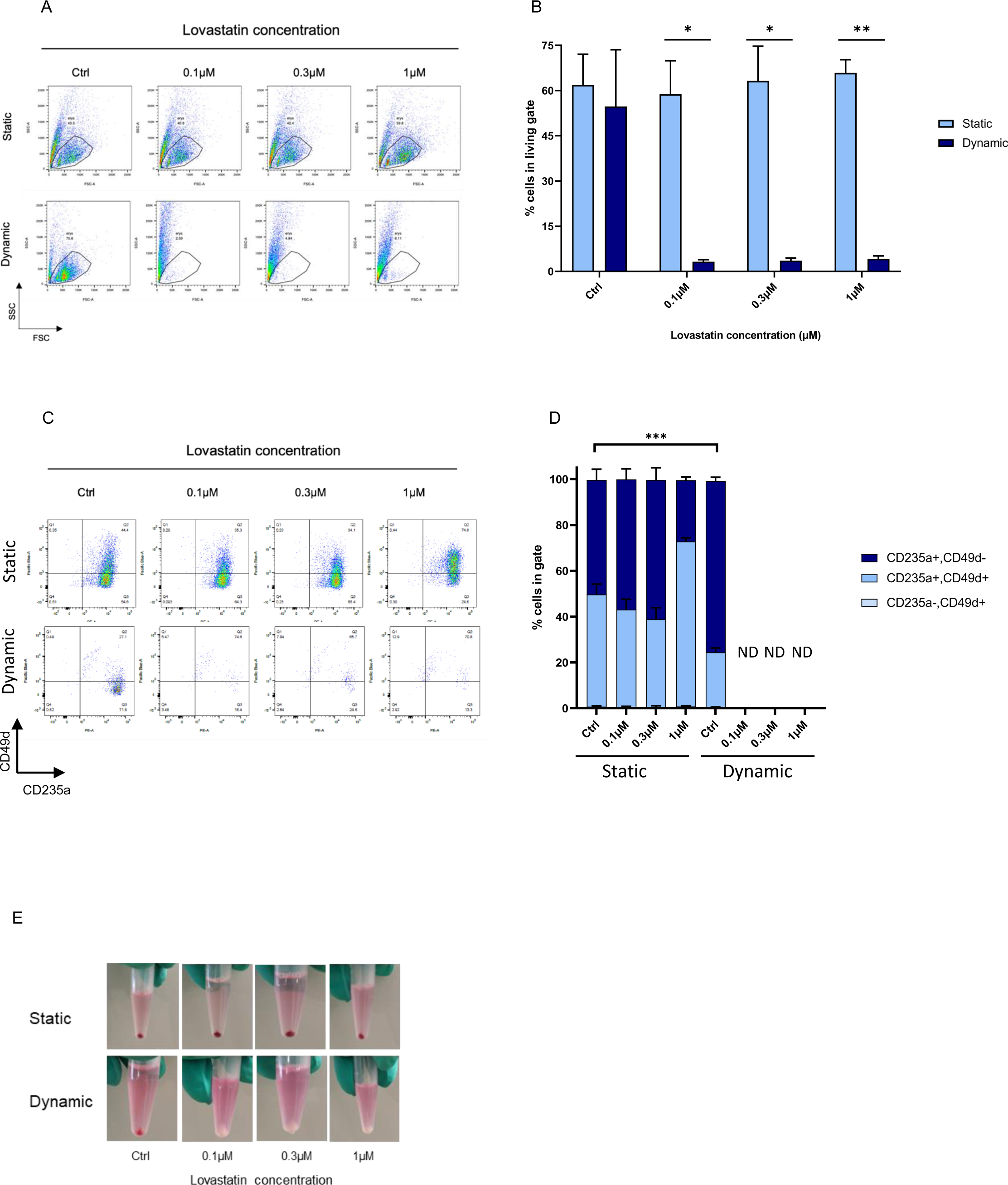
Lovastatin inhibition of HMGCR prevents cells adaptation to shear stress environment. Erythroblasts (EBL) of three different donors (n=3) were incubated with different concentrations of Lovastatin on day 0 of differentiation and cultured in static and dynamic conditions. Erythroid cells were harvested on day 12 of differentiation and analysed by flow cytometry. A) Representative example of living gate strategy plotting forward scatter (FSC x-axis) vs side scatter (SSC y-axis) in cells differentiated in static and dynamic conditions treated with different concentrations of lovastatin. B) Quantified percentage of cells in the living gate, averaged for 3 donors (n=3). C) Maturation states of cells differentiated in static and dynamic conditions incubated with different concentrations of lovastatin according to anti-CD49d (Alexa Fluor 405, y-axis) and anti-CD235a (PE, x-axis) staining. D) Quantified maturation states of cells differentiate in static and dynamic incubated with different concentrations of lovastatin were measured as described in Figure 1A. Percentage of cells in gates was averaged for 3 donors (n=3). ND indicates not-detectable measurement due to the low percentage of cells in the living gate. E) Representative picture of pelleted cells on day 12 of differentiation incubated with different concentration of lovastatin. B and D) 2way Anova test was performed, with *** p<0.001 ** p<0.01 and *p<0.05. Unless marked, no significance was observed.

We were unable to characterize differentiation markers for dynamic cultures in presence of lovastatin because most of the cells did not survive the HMGCR inhibition (Figure 6C, D). Overall, the data shows that HMGCR enzymatic inhibition was not tolerated during EBL differentiation under turbulent culture conditions and thus it is required for the adaptive response of EBL to orbital shaking.

## Discussion

EBL can be expanded to large numbers, sufficient to produce transfusion products^4^ (for review, see Di buduo^38^). However, cost-effective upscaling is crucial for the implementation of such products. To this end, advances in high cellular density cultures, media components and stirred tank bioreactor perfusion systems are required. As most upscaling systems introduce mixing by stirring, shaking and/or sparging of air/oxygen, the effects of mechanical stress on erythroid cells need to be evaluated.

Here we show that shear stress during CD49d^+^ CD235^low^ EBL maturation to enucleated CD49^-^ CD235^high^ reticulocytes accelerates erythroid differentiation with no effects on cell viability. We used a maximum shear stress of 1.4 Pa through orbital shaking at 0.7g, mimicking the shear stress experienced at the impeller of our stirred-tank-reactors^4^ and based on Odeleye^33^. Orbital shaking altered the EBL transcriptional program, which indicated that erythroid cells adapt to turbulent environments. The highest effect of mechanical forces on the acceleration of cell differentiation was detected within the first 4 days of erythroid differentiation.

Continuous orbital shaking (dynamic condition) changed the EBL transcriptional program. Interestingly, similar transcriptional responses to shear stress were also observed in other cell-types (*e.g.* 293 freestyle and endothelial cells).^39,40^ Characterization of the RNA expression throughout the first 4 days of differentiation revealed >500 DEGs between dynamic and static cultures. Among biological processes upregulated in dynamic cultures, the cholesterol biosynthesis pathway was highly enriched. This process may represent one of the aforementioned adaptation processes, enabling cells to withstand the shear stress experienced, ensuring a more rigid membrane by the increase of plasma membrane cholesterol content.

Mechanistically, we found several members of the cholesterol biosynthesis pathways being upregulated in dynamic conditions, such as SQLE, HMGCS1 and HMGCR. The latter is a key rate limiting catalyst in this process^41^ (for review, see Sharpe^42^). HMGCR is a target gene of Steroid Responsive Element Binding Proteins (SREBPs).^43^ Some SREBP regulating proteins such as HSD17B7, AACS and INSIG1 (for review, see Dong^44^) are also upregulated in dynamic conditions. Specifically, INSIG1 is involved in the transport of SREBP from the ER to the Golgi where it will be cleaved and activated.

Increased HMGCR expression correlates with higher cholesterol concentration in cells differentiated in dynamic compared to static conditions. This was associated with higher stability of reticulocytes cultured in dynamic conditions, comparable to RBC control. In contrast, cells differentiated in static showed lower stability, compared to both reticulocytes differentiated in shaker and RBC control. These data suggest that erythroid cells are adapting to shear stress by changing plasma-membrane fluidity (PMF) through the production of cholesterol rich membranes able to withstand shear.

Notably, HMGCR converts HMG-Co-enzyme A into mevalonic acid, a known pharmacological target of cholesterol decreasing therapeutics like statins.^36^ We then differentiated EBL in dynamic conditions in the presence of Lovastatin. The evidence that cells differentiated in dynamic conditions do not survive to the inhibition of HMGCR confirms how the up-regulation of genes involved in the cholesterol biosynthetic pathway represents an adaptation mechanism of the cell to shear stress environments.

Indeed, it has recently been shown that cholesterol supplementation and incorporation in the plasma membrane during erythroid differentiation is necessary to allow cultured erythrocytes to withstand osmotic stress comparable to native erythrocytes.^34^ It will be important to assess if the differences in plasma membrane composition of reticulocytes obtained in orbital shaker cultures have effects on the plasma membrane functionality. Indeed, it is known that high membrane cholesterol affects RBC O2 and CO2 transport and delivery.^45^ Moreover, more rigid membranes could have a negative effect on mechanosensing, which is dependent on the membrane mechanical properties and organisation of membrane cholesterol domains to coordinate the activity of mechano-channels.^46^ We and others have reported that mechanosensing proteins are present and functional in the erythroid membrane.^13,15,16^ EBL differentiate on central macrophages in erythroid islands in the bone marrow, where they may experience shear forces through crawling and dynamic contacts with these central macrophages.^47,48^ There may be an unknown but specific role for mechanosensors of hematopoietic progenitors and EBL during migration, sensing of tissue stiffness or in cell-cell contacts. Investigation of the putative mechanisms that regulate cell adaptation to mechanical forces, could be useful to understand how cells regulate their transcriptional program upon shear stress also in the bone marrow environment.

In conclusion, the data presented here clearly show that erythroid precursors are able to withstand and adapt to significant shear stress during differentiation to enucleated reticulocytes. Shear stress does not affect cell viability but accelerates the differentiation process. We have uncovered that cholesterol metabolism is essential for shear stress adaptation. The accelerated differentiation program allows a faster production protocol leading to a more cost-effective process and possibly permitting the upscale of EBL differentiation protocols into stirred-bioreactor setups.

