## Supplemental data for "Differentiating erythroblasts adapt to turbulent flow by accelerating maturation and activating cholesterol biosynthesis"

**Supplemental table 1. Genes differently expressed in static and dynamic condition on day 1 of differentiation**

| ensembl_gene_id | external_gene_name | logCPM | logFC | FDR |
|---|---|---|---|---|
| ENSG00000125740 | FOSB | 5.938769 | -2.09251 | 1.88E-15 |
| ENSG00000170345 | FOS | 8.004931 | -1.35395 | 1.25E-05 |
| ENSG00000167995 | BEST1 | 3.775391 | 1.751376 | 3.5E-05 |
| ENSG00000019582 | CD74 | 4.800891 | -1.20418 | 0.001224 |
| ENSG00000177675 | CD163L1 | 1.408855 | -2.60393 | 0.004568 |
| ENSG00000169429 | CXCL8 | 1.854446 | -2.28759 | 0.006452 |
| ENSG00000118785 | SPP1 | 2.496788 | -1.77215 | 0.041089 |
| ENSG00000038945 | MSR1 | 2.37929 | -1.8057 | 0.041089 |

**Supplemental table 2. Genes differently expressed in static and dynamic condition on day 2 of differentiation**

| ensembl_gene_id | external_gene_name | logCPM | logFC | FDR |
|---|---|---|---|---|
| ENSG00000125740 | FOSB | 5.9387691 | -2.34392 | 2.48E-19 |
| ENSG00000204389 | HSPA1A | 6.967433 | 1.473755 | 3.01E-07 |
| ENSG00000165507 | DEPP1 | 4.2197878 | -1.51115 | 0.000139 |
| ENSG00000204388 | HSPA1B | 7.6078625 | 1.189943 | 0.000338 |
| ENSG00000174358 | SLC6A19 | 6.0558441 | 1.168659 | 0.000454 |
| ENSG00000170345 | FOS | 8.0049305 | -1.15735 | 0.000548 |
| ENSG00000167244 | IGF2 | 6.3080995 | 1.041252 | 0.005299 |
| ENSG00000038945 | MSR1 | 2.3792899 | -2.01117 | 0.00877 |
| ENSG00000004399 | PLXND1 | 4.8822408 | -0.99182 | 0.03553 |
| ENSG00000101255 | TRIB3 | 6.9234316 | 0.937038 | 0.03553 |
| ENSG00000129538 | RNASE1 | 3.2076914 | -1.37583 | 0.046173 |
| ENSG00000197629 | MPEG1 | 1.5731138 | -2.13043 | 0.046173 |

**Supplemental table 3. Genes differently expressed in static and dynamic condition on day 4 of differentiation**

| ensembl_gene_id | external_gene_name | logCPM | logFC | FDR |
|---|---|---|---|---|
| ENSG00000224186 | PITX1-AS1 | 0.533431 | -3.39096 | 3.73E-05 |
| ENSG00000224356 | | -0.203058 | -3.18355 | 0.017156 |
| ENSG00000187566 | NHLRC1 | -0.297141 | -3.07042 | 0.025647 |
| ENSG00000101115 | SALL4 | 0.170786 | -2.97715 | 0.035668 |
| ENSG00000250072 | SH3TC2-DT | 0.483013 | -2.86322 | 0.010878 |
| ENSG00000267886 | | 0.760141 | -2.82728 | 0.007285 |
| ENSG00000172508 | CARNS1 | 2.66465 | -2.77117 | 1.1E-07 |
| ENSG00000232653 | GOLGA8N | 0.10616 | -2.71655 | 0.004813 |
| ENSG00000183828 | NUDT14 | 0.341323 | -2.61545 | 0.004464 |
| ENSG00000236548 | RNF217-AS1 | 0.508542 | -2.53557 | 0.005441 |

| | | | | |
|---|---|---|---|---|
| ENSG00000244398 | RPL36AP37 | -0.100642 | -2.48967 | 0.042593 |
| ENSG00000103740 | ACSBG1 | 5.699957 | -2.479 | 3.65E-18 |
| ENSG00000069011 | PITX1 | 2.524508 | -2.43249 | 1.44E-05 |
| ENSG00000199377 | RNU5F-1 | 1.596615 | -2.42146 | 0.008297 |
| ENSG00000049283 | EPN3 | 0.705265 | -2.34557 | 0.035668 |
| ENSG00000125246 | CLYBL | 0.820648 | -2.29001 | 0.021314 |
| ENSG00000214049 | UCA1 | 2.532919 | -2.27764 | 0.003654 |
| ENSG00000104140 | RHOV | 0.49822 | -2.25515 | 0.016684 |
| ENSG00000171401 | KRT13 | 1.4278 | -2.24111 | 0.040228 |
| ENSG00000173988 | LRRC63 | 0.236884 | -2.1971 | 0.035276 |
| ENSG00000141314 | RHBDL3 | 0.23548 | -2.18118 | 0.019225 |
| ENSG00000184221 | OLIG1 | 1.750435 | -2.16635 | 0.006181 |
| ENSG00000243323 | PTPRVP | 1.124297 | -2.15901 | 0.024816 |
| ENSG00000162551 | ALPL | 1.876187 | -2.13642 | 0.00545 |
| ENSG00000222724 | RNU2-63P | 2.213942 | -2.13573 | 0.002085 |
| ENSG00000196188 | CTSE | 2.606214 | -2.13227 | 0.000359 |
| ENSG00000077935 | SMC1B | 0.464059 | -2.13211 | 0.008444 |
| ENSG00000000003 | TSPAN6 | 0.503843 | -2.11219 | 0.035276 |
| ENSG00000223855 | PDGFA-DT | 0.170719 | -2.07946 | 0.034192 |
| ENSG00000229005 | HNF4A-AS1 | 0.72671 | -2.06852 | 0.019523 |
| ENSG00000167701 | GPT | 2.375136 | -1.99231 | 0.012717 |
| ENSG00000149596 | JPH2 | 1.063884 | -1.95998 | 0.008805 |
| ENSG00000168502 | MTCL1 | 1.769815 | -1.94822 | 0.015328 |
| ENSG00000143382 | ADAMTSL4 | 1.461802 | -1.90024 | 0.012717 |
| ENSG00000166689 | PLEKHA7 | 1.559811 | -1.89371 | 0.02522 |
| ENSG00000164920 | OSR2 | 3.355933 | -1.89353 | 2.11E-06 |
| ENSG00000134539 | KLRD1 | 2.256219 | -1.89003 | 0.001712 |
| ENSG00000196660 | SLC30A10 | 3.930601 | -1.88056 | 1.12E-07 |
| ENSG00000242715 | CCDC169 | 1.976669 | -1.87358 | 0.038412 |
| ENSG00000166446 | CDYL2 | 2.948108 | -1.75956 | 0.000841 |
| ENSG00000112667 | DNPH1 | 2.43134 | -1.71206 | 0.012051 |
| ENSG00000257365 | FNTB | 2.172696 | -1.69699 | 0.018017 |
| ENSG00000164051 | CCDC51 | 2.438417 | -1.68847 | 0.018213 |
| ENSG00000188191 | PRKAR1B | 3.396162 | -1.66471 | 0.001306 |
| ENSG00000187997 | C17orf99 | 3.533449 | -1.66337 | 3.94E-05 |
| ENSG00000112425 | EPM2A | 2.557437 | -1.65336 | 0.003615 |
| ENSG00000073536 | NLE1 | 2.804229 | -1.64514 | 0.019225 |
| ENSG00000157870 | PRXL2B | 3.565884 | -1.61545 | 0.00049 |
| ENSG00000204815 | ODAD4 | 2.274256 | -1.60757 | 0.013781 |
| ENSG00000169247 | SH3TC2 | 4.220759 | -1.59502 | 2.34E-05 |
| ENSG00000270977 | | 1.71115 | -1.58792 | 0.041463 |
| ENSG00000078900 | TP73 | 2.458843 | -1.58006 | 0.014121 |
| ENSG00000164879 | CA3 | 2.481803 | -1.55503 | 0.008982 |
| ENSG00000165507 | DEPP1 | 4.219788 | -1.52383 | 3.2E-06 |
| ENSG00000135052 | GOLM1 | 4.857131 | -1.49661 | 1.78E-06 |

| | | | | |
|---|---|---|---|---|
| ENSG00000162062 | TEDC2 | 3.109525 | -1.47233 | 0.007225 |
| ENSG00000258738 | BAZ1A-AS1 | 2.174222 | -1.45922 | 0.042773 |
| ENSG00000132016 | BRME1 | 3.161283 | -1.45705 | 0.002398 |
| ENSG00000125520 | SLC2A4RG | 4.466139 | -1.45069 | 8.14E-06 |
| ENSG00000184083 | FAM120C | 2.813366 | -1.42543 | 0.025647 |
| ENSG00000140280 | LYSMD2 | 3.803499 | -1.41919 | 0.000355 |
| ENSG00000173801 | JUP | 3.14952 | -1.4158 | 0.008196 |
| ENSG00000144354 | CDCA7 | 3.408654 | -1.41307 | 0.007163 |
| ENSG00000152454 | ZNF256 | 2.941406 | -1.40759 | 0.015795 |
| ENSG00000184208 | C22orf46 | 3.742779 | -1.40749 | 0.000673 |
| ENSG00000095303 | PTGS1 | 3.569782 | -1.40446 | 0.000798 |
| ENSG00000280434 | | 4.001962 | -1.40399 | 0.000328 |
| ENSG00000162496 | DHRS3 | 3.707645 | -1.36133 | 0.001012 |
| ENSG00000163872 | YEATS2 | 5.059178 | -1.35119 | 5.78E-06 |
| ENSG00000161267 | BDH1 | 4.023925 | -1.34718 | 0.0011 |
| ENSG00000064201 | TSPAN32 | 5.3842 | -1.34689 | 8.16E-06 |
| ENSG00000100346 | CACNA1I | 3.227438 | -1.34605 | 0.005319 |
| ENSG00000224259 | LINC01133 | 4.358994 | -1.34267 | 0.000121 |
| ENSG00000117016 | RIMS3 | 4.455774 | -1.3413 | 0.000245 |
| ENSG00000156966 | B3GNT7 | 2.592923 | -1.34035 | 0.035668 |
| ENSG00000117013 | KCNQ4 | 2.492201 | -1.33492 | 0.047336 |
| ENSG00000279175 | | 3.396276 | -1.33116 | 0.006705 |
| ENSG00000162073 | PAQR4 | 4.026221 | -1.31762 | 0.000673 |
| ENSG00000249684 | | 2.636011 | -1.30204 | 0.043736 |
| ENSG00000167880 | EVPL | 2.930155 | -1.29846 | 0.019682 |
| ENSG00000085117 | CD82 | 5.630507 | -1.29601 | 1.29E-05 |
| ENSG00000154920 | EME1 | 3.537143 | -1.28737 | 0.00545 |
| ENSG00000168961 | LGALS9 | 3.16225 | -1.28386 | 0.009627 |
| ENSG00000182378 | PLCXD1 | 5.233371 | -1.28203 | 3.36E-05 |
| ENSG00000174306 | ZHX3 | 5.081738 | -1.28053 | 5.29E-05 |
| ENSG00000162065 | TBC1D24 | 4.508956 | -1.26193 | 0.000554 |
| ENSG00000143627 | PKLR | 6.42144 | -1.24789 | 2.34E-05 |
| ENSG00000188157 | AGRN | 4.691211 | -1.2412 | 0.000182 |
| ENSG00000257800 | FNBP1P1 | 2.879118 | -1.23644 | 0.034192 |
| ENSG00000124610 | H1-1 | 3.391921 | -1.2173 | 0.018213 |
| ENSG00000112297 | CRYBG1 | 3.577258 | -1.21492 | 0.012332 |
| ENSG00000279602 | | 3.636921 | -1.2092 | 0.003654 |
| ENSG00000122694 | GLIPR2 | 4.489396 | -1.20491 | 0.000388 |
| ENSG00000037042 | TUBG2 | 4.772938 | -1.19222 | 0.000393 |
| ENSG00000212195 | U3 | 3.178293 | -1.19084 | 0.03393 |
| ENSG00000156509 | FBXO43 | 3.016618 | -1.18195 | 0.029596 |
| ENSG00000187792 | ZNF70 | 3.351057 | -1.1723 | 0.021267 |
| ENSG00000226688 | ENTPD1-AS1 | 3.841434 | -1.16745 | 0.005783 |
| ENSG00000105202 | FBL | 5.956177 | -1.15854 | 0.000115 |
| ENSG00000177432 | NAP1L5 | 3.6561 | -1.15077 | 0.007864 |
| ENSG00000182054 | IDH2 | 7.382393 | -1.14542 | 7.8E-05 |

| | | | | |
|---|---|---|---|---|
| ENSG00000007944 | MYLIP | 5.511232 | -1.14125 | 9.68E-05 |
| ENSG00000126001 | CEP250 | 6.212315 | -1.14109 | 0.000113 |
| ENSG00000196535 | MYO18A | 5.806364 | -1.14098 | 0.000132 |
| ENSG00000162924 | REL | 5.263331 | -1.11777 | 0.000212 |
| ENSG00000176912 | TYMSOS | 2.980241 | -1.11767 | 0.046999 |
| ENSG00000276043 | UHRF1 | 6.690778 | -1.11612 | 0.000139 |
| ENSG00000274210 | RNVU1-27 | 4.067272 | -1.11373 | 0.004237 |
| ENSG00000077514 | POLD3 | 5.566877 | -1.10923 | 0.000355 |
| ENSG00000276180 | H4C9 | 8.370066 | -1.0997 | 0.000254 |
| ENSG00000086289 | EPDR1 | 5.284486 | -1.09678 | 0.000554 |
| ENSG00000188677 | PARVB | 5.611488 | -1.09495 | 0.00048 |
| ENSG00000156398 | SFXN2 | 4.006493 | -1.09173 | 0.023532 |
| ENSG00000176974 | SHMT1 | 4.627141 | -1.09151 | 0.002085 |
| ENSG00000141577 | CEP131 | 4.338131 | -1.08573 | 0.002872 |
| ENSG00000131127 | ZNF141 | 4.1967 | -1.08533 | 0.005786 |
| ENSG00000110665 | C11orf21 | 4.570552 | -1.0817 | 0.004489 |
| ENSG00000146281 | PM20D2 | 3.421139 | -1.07827 | 0.023064 |
| ENSG00000175322 | ZNF519 | 3.620916 | -1.07028 | 0.034995 |
| ENSG00000148154 | UGCG | 5.457286 | -1.06819 | 0.000435 |
| ENSG00000112759 | SLC29A1 | 6.594477 | -1.06541 | 0.000357 |
| ENSG00000167895 | TMC8 | 4.296822 | -1.06453 | 0.006925 |
| ENSG00000167363 | FN3K | 5.7507 | -1.06167 | 0.00049 |
| ENSG00000276410 | H2BC3 | 8.30962 | -1.06118 | 0.00044 |
| ENSG00000105173 | CCNE1 | 5.698685 | -1.06096 | 0.000588 |
| ENSG00000146540 | C7orf50 | 4.41528 | -1.05403 | 0.005089 |
| ENSG00000164402 | SEPTIN8 | 5.975788 | -1.05168 | 0.000552 |
| ENSG00000111261 | MANSC1 | 3.409365 | -1.05011 | 0.035668 |
| ENSG00000128973 | CLN6 | 4.84831 | -1.04817 | 0.002195 |
| ENSG00000128872 | TMOD2 | 5.204397 | -1.04721 | 0.000813 |
| ENSG00000111843 | TMEM14C | 7.042396 | -1.04629 | 0.000396 |
| ENSG00000181938 | GINS3 | 4.32172 | -1.04586 | 0.005734 |
| ENSG00000131153 | GINS2 | 5.091016 | -1.04448 | 0.00188 |
| ENSG00000073111 | MCM2 | 7.53374 | -1.03164 | 0.00056 |
| ENSG00000063761 | ADCK1 | 3.888007 | -1.0315 | 0.014836 |
| ENSG00000107201 | RIGI | 3.684442 | -1.02599 | 0.016684 |
| ENSG00000117298 | ECE1 | 5.459849 | -1.02271 | 0.001133 |
| ENSG00000084444 | FAM234B | 4.393279 | -1.01832 | 0.006661 |
| ENSG00000198208 | RPS6KL1 | 4.644627 | -1.01748 | 0.017527 |
| ENSG00000144401 | METTL21A | 3.643368 | -1.01732 | 0.040997 |
| ENSG00000177842 | ZNF620 | 4.207961 | -1.0154 | 0.012528 |
| ENSG00000178921 | PFAS | 5.239793 | -1.01269 | 0.006925 |
| ENSG00000215883 | CYB5RL | 3.813153 | -1.0105 | 0.030349 |
| ENSG00000232931 | LINC00342 | 3.573894 | -1.0093 | 0.037563 |
| ENSG00000187244 | BCAM | 5.93737 | -1.00121 | 0.000879 |
| ENSG00000125148 | MT2A | 3.886727 | -0.99997 | 0.016237 |
| ENSG00000235194 | PPP1R3E | 3.407164 | -0.99693 | 0.045955 |

| | | | | |
|---|---|---|---|---|
| ENSG00000197153 | H3C12 | 8.008332 | -0.99563 | 0.0011 |
| ENSG00000138796 | HADH | 3.944032 | -0.98829 | 0.028078 |
| ENSG00000070526 | ST6GALNAC1 | 3.834944 | -0.98189 | 0.02936 |
| ENSG00000075340 | ADD2 | 9.221962 | -0.97935 | 0.00186 |
| ENSG00000214189 | ZNF788P | 4.699701 | -0.97561 | 0.006925 |
| ENSG00000129244 | ATP1B2 | 7.207298 | -0.97456 | 0.001102 |
| ENSG00000100297 | MCM5 | 7.8014 | -0.96846 | 0.001518 |
| ENSG00000097021 | ACOT7 | 4.064815 | -0.96001 | 0.029093 |
| ENSG00000164649 | CDCA7L | 4.324996 | -0.95926 | 0.016515 |
| ENSG00000100350 | FOXRED2 | 5.112603 | -0.95802 | 0.004944 |
| ENSG00000118503 | TNFAIP3 | 3.4101 | -0.95262 | 0.049328 |
| ENSG00000257167 | TMPO-AS1 | 3.956597 | -0.94797 | 0.023064 |
| ENSG00000160446 | ZDHHC12 | 3.982417 | -0.94576 | 0.02522 |
| ENSG00000261236 | BOP1 | 5.071997 | -0.94537 | 0.009021 |
| ENSG00000146373 | RNF217 | 5.286712 | -0.94461 | 0.003658 |
| ENSG00000198521 | ZNF43 | 3.767087 | -0.94173 | 0.040694 |
| ENSG00000188735 | TMEM120B | 6.184462 | -0.93998 | 0.002398 |
| ENSG00000179981 | TSHZ1 | 3.829422 | -0.93676 | 0.030804 |
| ENSG00000275835 | TUBGCP5 | 4.050156 | -0.9293 | 0.030804 |
| ENSG00000163170 | BOLA3 | 4.31253 | -0.92767 | 0.027995 |
| ENSG00000115687 | PASK | 5.419363 | -0.92739 | 0.004944 |
| ENSG00000145990 | GFOD1 | 5.904878 | -0.92573 | 0.003105 |
| ENSG00000104738 | MCM4 | 8.122153 | -0.92444 | 0.003433 |
| ENSG00000076003 | MCM6 | 6.971177 | -0.9202 | 0.003095 |
| ENSG00000211459 | MT-RNR1 | 7.692087 | -0.9179 | 0.003095 |
| ENSG00000162337 | LRP5 | 4.554913 | -0.91201 | 0.014388 |
| ENSG00000111684 | LPCAT3 | 6.35938 | -0.90616 | 0.004036 |
| ENSG00000068137 | PLEKHH3 | 6.695805 | -0.90571 | 0.003644 |
| ENSG00000126215 | XRCC3 | 5.551926 | -0.904 | 0.006205 |
| ENSG00000112118 | MCM3 | 7.024729 | -0.9029 | 0.003684 |
| ENSG00000163864 | NMNAT3 | 4.717102 | -0.90119 | 0.020219 |
| ENSG00000167792 | NDUFV1 | 5.450453 | -0.90081 | 0.007285 |
| ENSG00000181350 | LRRC75A | 5.369798 | -0.89897 | 0.005736 |
| ENSG00000214076 | CPSF1P1 | 4.589916 | -0.89747 | 0.015845 |
| ENSG00000116857 | TMEM9 | 3.996958 | -0.89535 | 0.036488 |
| ENSG00000094804 | CDC6 | 6.184943 | -0.89167 | 0.00545 |
| ENSG00000177595 | PIDD1 | 4.675306 | -0.89148 | 0.01527 |
| ENSG00000166803 | PCLAF | 4.756186 | -0.89067 | 0.014121 |
| ENSG00000100336 | APOL4 | 4.579073 | -0.88997 | 0.036488 |
| ENSG00000141456 | PELP1 | 4.984863 | -0.88948 | 0.014689 |
| ENSG00000166025 | AMOTL1 | 5.038114 | -0.88948 | 0.009324 |
| ENSG00000240342 | RPS2P5 | 5.894781 | -0.8894 | 0.006705 |
| ENSG00000273703 | H2BC14 | 8.488118 | -0.88767 | 0.005812 |
| ENSG00000155980 | KIF5A | 4.6641 | -0.88574 | 0.016169 |
| ENSG00000109265 | CRACD | 5.045185 | -0.88307 | 0.01442 |
| ENSG00000112576 | CCND3 | 7.478498 | -0.88257 | 0.00518 |

| | | | | |
|---|---|---|---|---|
| ENSG00000273542 | H4C12 | 3.959556 | -0.88236 | 0.042773 |
| ENSG00000273802 | H2BC8 | 8.831752 | -0.87731 | 0.007065 |
| ENSG00000059573 | ALDH18A1 | 5.241185 | -0.87535 | 0.019225 |
| ENSG00000152240 | HAUS1 | 4.506775 | -0.87402 | 0.025647 |
| ENSG00000171858 | RPS21 | 7.420296 | -0.87258 | 0.005783 |
| ENSG00000072786 | STK10 | 5.639786 | -0.87227 | 0.007441 |
| ENSG00000133488 | SEC14L4 | 6.227975 | -0.87171 | 0.006208 |
| ENSG00000004809 | SLC22A16 | 5.990543 | -0.87129 | 0.006412 |
| ENSG00000240583 | AQP1 | 7.314263 | -0.86982 | 0.005517 |
| ENSG00000083857 | FAT1 | 4.715265 | -0.86676 | 0.014341 |
| ENSG00000141873 | SLC39A3 | 4.693859 | -0.86651 | 0.021314 |
| ENSG00000146729 | NIPSNAP2 | 4.187462 | -0.86408 | 0.040162 |
| ENSG00000276903 | H2AC16 | 8.505621 | -0.86375 | 0.00766 |
| ENSG00000141524 | TMC6 | 6.552241 | -0.86288 | 0.007285 |
| ENSG00000249115 | HAUS5 | 4.863382 | -0.862 | 0.016684 |
| ENSG00000067533 | RRP15 | 4.858325 | -0.86171 | 0.017653 |
| ENSG00000275221 | H2AC15 | 6.633203 | -0.85983 | 0.00656 |
| ENSG00000143476 | DTL | 6.897134 | -0.85952 | 0.006661 |
| ENSG00000168405 | CMAHP | 4.861581 | -0.85445 | 0.019225 |
| ENSG00000141905 | NFIC | 5.363067 | -0.85364 | 0.010755 |
| ENSG00000186767 | SPIN4 | 5.252541 | -0.85259 | 0.012051 |
| ENSG00000198752 | CDC42BPB | 5.065935 | -0.85066 | 0.013516 |
| ENSG00000127564 | PKMYT1 | 6.328047 | -0.84898 | 0.007864 |
| ENSG00000179750 | APOBEC3B | 4.243874 | -0.84819 | 0.043404 |
| ENSG00000205213 | LGR4 | 5.012461 | -0.84747 | 0.016515 |
| ENSG00000106991 | ENG | 5.891819 | -0.84719 | 0.015845 |
| ENSG00000157985 | AGAP1 | 4.460588 | -0.84222 | 0.029635 |
| ENSG00000152229 | PSTPIP2 | 4.591988 | -0.8282 | 0.037564 |
| ENSG00000143537 | ADAM15 | 4.455854 | -0.82776 | 0.032557 |
| ENSG00000124635 | H2BC11 | 8.600436 | -0.8254 | 0.013199 |
| ENSG00000196584 | XRCC2 | 5.499205 | -0.82404 | 0.016515 |
| ENSG00000111716 | LDHB | 6.796984 | -0.81906 | 0.01527 |
| ENSG00000280184 | | 5.333169 | -0.81552 | 0.014174 |
| ENSG00000149260 | CAPN5 | 4.218156 | -0.81539 | 0.045693 |
| ENSG00000277075 | H2AC8 | 9.451935 | -0.81534 | 0.016684 |
| ENSG00000116120 | FARSB | 5.939503 | -0.81331 | 0.017653 |
| ENSG00000213445 | SIPA1 | 5.337279 | -0.81099 | 0.016694 |
| ENSG00000276966 | H4C5 | 9.244041 | -0.80862 | 0.017156 |
| ENSG00000168056 | LTBP3 | 4.29542 | -0.8086 | 0.044371 |
| ENSG00000165244 | ZNF367 | 6.240871 | -0.80821 | 0.014243 |
| ENSG00000100714 | MTHFD1 | 7.015288 | -0.8052 | 0.014632 |
| ENSG00000197785 | ATAD3A | 4.770098 | -0.80515 | 0.040186 |
| ENSG00000099901 | RANBP1 | 6.367184 | -0.80507 | 0.016823 |
| ENSG00000167747 | C19orf48 | 5.64935 | -0.80445 | 0.019388 |
| ENSG00000170629 | DPY19L2P2 | 5.469806 | -0.80149 | 0.018349 |
| ENSG00000147536 | GINS4 | 4.865945 | -0.79881 | 0.033017 |

| | | | | |
|---|---|---|---|---|
| ENSG00000129355 | CDKN2D | 5.19681 | -0.79872 | 0.01838 |
| ENSG00000132182 | NUP210 | 7.550044 | -0.79776 | 0.015845 |
| ENSG00000198755 | RPL10A | 7.6772 | -0.79761 | 0.016684 |
| ENSG00000103202 | NME4 | 6.606463 | -0.79421 | 0.016515 |
| ENSG00000160957 | RECQL4 | 6.081448 | -0.79332 | 0.017813 |
| ENSG00000198039 | ZNF273 | 5.456014 | -0.79305 | 0.019388 |
| ENSG00000138821 | SLC39A8 | 6.326903 | -0.79182 | 0.019225 |
| ENSG00000128274 | A4GALT | 6.278151 | -0.79145 | 0.01874 |
| ENSG00000101220 | C20orf27 | 4.842259 | -0.79027 | 0.036128 |
| ENSG00000143493 | INTS7 | 5.558877 | -0.78808 | 0.022641 |
| ENSG00000128951 | DUT | 6.602324 | -0.78716 | 0.017509 |
| ENSG00000169891 | REPS2 | 5.822559 | -0.78551 | 0.026639 |
| ENSG00000146143 | PRIM2 | 4.996375 | -0.78392 | 0.032099 |
| ENSG00000154518 | ATP5MC3 | 5.671477 | -0.78241 | 0.027293 |
| ENSG00000119471 | HSDL2 | 5.323833 | -0.78196 | 0.026639 |
| ENSG00000210082 | MT-RNR2 | 8.676453 | -0.78071 | 0.021314 |
| ENSG00000108219 | TSPAN14 | 6.839363 | -0.78064 | 0.017276 |
| ENSG00000274997 | H2AC12 | 9.050281 | -0.77751 | 0.023534 |
| ENSG00000171793 | CTPS1 | 5.676051 | -0.77726 | 0.026972 |
| ENSG00000173848 | NET1 | 6.774105 | -0.77643 | 0.017276 |
| ENSG00000196230 | TUBB | 10.53651 | -0.77299 | 0.029846 |
| ENSG00000171848 | RRM2 | 9.444067 | -0.77194 | 0.026445 |
| ENSG00000171914 | TLN2 | 4.735512 | -0.76999 | 0.040538 |
| ENSG00000164045 | CDC25A | 6.236076 | -0.76941 | 0.022604 |
| ENSG00000198721 | ECI2 | 4.847409 | -0.76837 | 0.044284 |
| ENSG00000113368 | LMNB1 | 6.631432 | -0.76762 | 0.021314 |
| ENSG00000186575 | NF2 | 6.368245 | -0.76729 | 0.023687 |
| ENSG00000197903 | H2BC12 | 9.7459 | -0.76503 | 0.029772 |
| ENSG00000106605 | BLVRA | 5.249122 | -0.7594 | 0.037563 |
| ENSG00000100316 | RPL3 | 9.205497 | -0.75857 | 0.030167 |
| ENSG00000277224 | H2BC7 | 9.359163 | -0.75727 | 0.030629 |
| ENSG00000141560 | FN3KRP | 4.829344 | -0.75446 | 0.045693 |
| ENSG00000100162 | CENPM | 5.053833 | -0.75423 | 0.036931 |
| ENSG00000066735 | KIF26A | 7.565027 | -0.75279 | 0.025196 |
| ENSG00000148218 | ALAD | 8.862022 | -0.75167 | 0.030572 |
| ENSG00000164442 | CITED2 | 9.203282 | -0.75075 | 0.031114 |
| ENSG00000156675 | RAB11FIP1 | 5.789284 | -0.74998 | 0.030282 |
| ENSG00000113569 | NUP155 | 6.77041 | -0.74887 | 0.02577 |
| ENSG00000100292 | HMOX1 | 6.597896 | -0.74874 | 0.023676 |
| ENSG00000198888 | MT-ND1 | 9.662039 | -0.74865 | 0.034697 |
| ENSG00000077152 | UBE2T | 5.35568 | -0.74646 | 0.034074 |
| ENSG00000275379 | H3C11 | 8.457203 | -0.74541 | 0.030804 |
| ENSG00000167088 | SNRPD1 | 5.655734 | -0.74494 | 0.037563 |
| ENSG00000110888 | CAPRIN2 | 6.982067 | -0.74456 | 0.025647 |
| ENSG00000136840 | ST6GALNAC4 | 6.612845 | -0.744 | 0.026782 |
| ENSG00000278705 | H4C2 | 8.477646 | -0.74376 | 0.031111 |

| | | | | |
|---|---|---|---|---|
| ENSG00000117676 | RPS6KA1 | 6.425891 | -0.74301 | 0.029205 |
| ENSG00000157734 | SNX22 | 7.161138 | -0.74227 | 0.025647 |
| ENSG00000184992 | BRI3BP | 6.028249 | -0.74145 | 0.034764 |
| ENSG00000116062 | MSH6 | 6.736933 | -0.74007 | 0.02936 |
| ENSG00000184678 | H2BC21 | 9.367388 | -0.73515 | 0.037276 |
| ENSG00000278637 | H4C1 | 8.186484 | -0.73501 | 0.033017 |
| ENSG00000275527 | | 6.45419 | -0.73489 | 0.029733 |
| ENSG00000085999 | RAD54L | 5.617527 | -0.7343 | 0.036082 |
| ENSG00000278272 | #N/A | 9.592124 | -0.73409 | 0.038881 |
| ENSG00000137210 | TMEM14B | 6.880654 | -0.73276 | 0.030706 |
| ENSG00000004777 | ARHGAP33 | 5.334763 | -0.73252 | 0.036931 |
| ENSG00000150712 | MTMR12 | 6.641937 | -0.7306 | 0.03111 |
| ENSG00000183032 | SLC25A21 | 5.265936 | -0.72962 | 0.045745 |
| ENSG00000231500 | RPS18 | 8.662302 | -0.72393 | 0.039413 |
| ENSG00000118971 | CCND2 | 5.737712 | -0.72389 | 0.042773 |
| ENSG00000073464 | CLCN4 | 6.585024 | -0.72265 | 0.036595 |
| ENSG00000140526 | ABHD2 | 6.774876 | -0.72136 | 0.033712 |
| ENSG00000067836 | ROGDI | 5.23026 | -0.72064 | 0.045693 |
| ENSG00000092853 | CLSPN | 7.055247 | -0.72005 | 0.032473 |
| ENSG00000196747 | H2AC13 | 9.62204 | -0.71856 | 0.045264 |
| ENSG00000164032 | H2AZ1 | 7.436622 | -0.71663 | 0.035668 |
| ENSG00000115107 | STEAP3 | 6.919963 | -0.71596 | 0.034764 |
| ENSG00000160949 | TONSL | 5.85131 | -0.71482 | 0.045264 |
| ENSG00000105810 | CDK6 | 6.524186 | -0.71265 | 0.044847 |
| ENSG00000062822 | POLD1 | 6.119512 | -0.70607 | 0.044847 |
| ENSG00000167513 | CDT1 | 6.609451 | -0.70581 | 0.040325 |
| ENSG00000100345 | MYH9 | 8.752863 | -0.7055 | 0.046934 |
| ENSG00000149269 | PAK1 | 6.832191 | -0.70452 | 0.040525 |
| ENSG00000048140 | TSPAN17 | 6.893931 | -0.70275 | 0.040778 |
| ENSG00000198728 | LDB1 | 7.466068 | -0.70256 | 0.042803 |
| ENSG00000113369 | ARRDC3 | 7.095343 | -0.70167 | 0.037248 |
| ENSG00000143799 | PARP1 | 6.666881 | -0.69922 | 0.045755 |
| ENSG00000123374 | CDK2 | 6.08572 | -0.69679 | 0.048775 |
| ENSG00000278463 | H2AC4 | 7.714457 | -0.69636 | 0.045264 |
| ENSG00000119969 | HELLS | 6.866046 | -0.69609 | 0.043404 |
| ENSG00000178999 | AURKB | 5.787458 | -0.69564 | 0.049683 |
| ENSG00000154760 | SLFN13 | 6.384579 | -0.69335 | 0.045693 |
| ENSG00000205542 | TMSB4X | 6.597143 | 0.677794 | 0.049983 |
| ENSG00000011198 | ABHD5 | 7.299966 | 0.691096 | 0.043404 |
| ENSG00000170348 | TMED10 | 6.401237 | 0.692225 | 0.045634 |
| ENSG00000130164 | LDLR | 7.275964 | 0.692429 | 0.043057 |
| ENSG00000213918 | DNASE1 | 6.117107 | 0.694861 | 0.044282 |
| ENSG00000196730 | DAPK1 | 8.073823 | 0.703691 | 0.043517 |
| ENSG00000121073 | SLC35B1 | 5.334922 | 0.703885 | 0.047336 |
| ENSG00000119285 | HEATR1 | 6.723802 | 0.704508 | 0.038412 |
| ENSG00000169155 | ZBTB43 | 5.87836 | 0.707385 | 0.040525 |

| | | | | |
|---|---|---|---|---|
| ENSG00000124333 | VAMP7 | 6.185621 | 0.711824 | 0.037185 |
| ENSG00000136235 | GPNMB | 6.12249 | 0.716709 | 0.035668 |
| ENSG00000180385 | EMC3-AS1 | 5.709843 | 0.719864 | 0.036488 |
| ENSG00000213090 | SCYL2P1 | 5.902735 | 0.721183 | 0.035394 |
| ENSG00000151065 | DCP1B | 5.134298 | 0.723818 | 0.041471 |
| ENSG00000101294 | HM13 | 5.420855 | 0.724068 | 0.040609 |
| ENSG00000072042 | RDH11 | 6.492418 | 0.725448 | 0.031114 |
| ENSG00000107959 | PITRM1 | 6.454126 | 0.730276 | 0.030804 |
| ENSG00000081181 | ARG2 | 6.859052 | 0.73049 | 0.028078 |
| ENSG00000197343 | ZNF655 | 5.359117 | 0.730781 | 0.036488 |
| ENSG00000179348 | GATA2 | 5.153084 | 0.734744 | 0.037185 |
| ENSG00000086061 | DNAJA1 | 7.848703 | 0.735068 | 0.030804 |
| ENSG00000067064 | IDI1 | 5.81424 | 0.739389 | 0.030631 |
| ENSG00000167996 | FTH1 | 12.3916 | 0.745632 | 0.043736 |
| ENSG00000163900 | TMEM41A | 5.100601 | 0.748174 | 0.034654 |
| ENSG00000135679 | MDM2 | 9.111007 | 0.752549 | 0.030706 |
| ENSG00000142657 | PGD | 7.129438 | 0.753162 | 0.022641 |
| ENSG00000110880 | CORO1C | 8.227258 | 0.758251 | 0.02577 |
| ENSG00000109906 | ZBTB16 | 5.521691 | 0.761038 | 0.023517 |
| ENSG00000155749 | FLACC1 | 7.05092 | 0.764109 | 0.018569 |
| ENSG00000197157 | SND1 | 6.820182 | 0.765294 | 0.019388 |
| ENSG00000263606 | CHORDC1P4 | 5.060456 | 0.769828 | 0.026972 |
| ENSG00000139793 | MBNL2 | 8.792066 | 0.772294 | 0.023687 |
| ENSG00000154803 | FLCN | 8.21946 | 0.775186 | 0.02113 |
| ENSG00000181026 | AEN | 7.415794 | 0.775737 | 0.017653 |
| ENSG00000112242 | E2F3 | 5.566791 | 0.779665 | 0.020848 |
| ENSG00000128016 | ZFP36 | 6.811313 | 0.796091 | 0.013677 |
| ENSG00000260641 | TSPAN5-DT | 5.266913 | 0.798586 | 0.017653 |
| ENSG00000140105 | WARS1 | 6.927169 | 0.798789 | 0.012975 |
| ENSG00000112343 | TRIM38 | 5.638656 | 0.802729 | 0.015226 |
| ENSG00000134265 | NAPG | 6.908906 | 0.803231 | 0.012332 |
| ENSG00000198814 | GK | 4.289182 | 0.805421 | 0.046594 |
| ENSG00000134324 | LPIN1 | 4.383317 | 0.80611 | 0.040186 |
| ENSG00000081760 | AACS | 5.38182 | 0.807071 | 0.016431 |
| ENSG00000138768 | USO1 | 4.954453 | 0.810707 | 0.021314 |
| ENSG00000186480 | INSIG1 | 7.351064 | 0.819528 | 0.010643 |
| ENSG00000164543 | STK17A | 7.977691 | 0.82116 | 0.01225 |
| ENSG00000113161 | HMGCR | 6.565818 | 0.821306 | 0.010878 |
| ENSG00000196182 | STK40 | 6.995087 | 0.823206 | 0.008868 |
| ENSG00000018189 | RUFY3 | 4.913326 | 0.829307 | 0.015845 |
| ENSG00000223509 | WHAMMP1 | 4.362078 | 0.832807 | 0.032099 |
| ENSG00000101255 | TRIB3 | 6.923432 | 0.835784 | 0.00756 |
| ENSG00000140450 | ARRDC4 | 4.691679 | 0.837947 | 0.019388 |
| ENSG00000107651 | SEC23IP | 5.927821 | 0.846648 | 0.008144 |
| ENSG00000151694 | ADAM17 | 5.779307 | 0.847136 | 0.008196 |
| ENSG00000074800 | ENO1 | 6.104731 | 0.849815 | 0.008299 |

| | | | | |
|---|---|---|---|---|
| ENSG00000047644 | WWC3 | 8.235561 | 0.859388 | 0.007569 |
| ENSG00000112972 | HMGCS1 | 7.828698 | 0.86073 | 0.007065 |
| ENSG00000164934 | DCAF13 | 4.655925 | 0.864928 | 0.016684 |
| ENSG00000099194 | SCD | 8.189264 | 0.867099 | 0.007171 |
| ENSG00000076685 | NT5C2 | 5.382739 | 0.86732 | 0.00749 |
| ENSG00000010404 | IDS | 3.963874 | 0.868299 | 0.037564 |
| ENSG00000157796 | WDR19 | 5.031004 | 0.880405 | 0.007855 |
| ENSG00000170876 | TMEM43 | 4.518012 | 0.884765 | 0.016684 |
| ENSG00000101236 | RNF24 | 7.303004 | 0.885931 | 0.004237 |
| ENSG00000163558 | PRKCI | 4.962116 | 0.888416 | 0.007679 |
| ENSG00000156206 | CFAP161 | 4.171633 | 0.902936 | 0.019388 |
| ENSG00000096384 | HSP90AB1 | 9.183807 | 0.906314 | 0.005254 |
| ENSG00000130827 | PLXNA3 | 3.892772 | 0.91538 | 0.030804 |
| ENSG00000163110 | PDLIM5 | 4.549904 | 0.920777 | 0.010693 |
| ENSG00000132256 | TRIM5 | 4.724969 | 0.924957 | 0.00756 |
| ENSG00000164506 | STXBP5 | 4.223866 | 0.929153 | 0.016745 |
| ENSG00000168439 | STIP1 | 7.094496 | 0.932267 | 0.002046 |
| ENSG00000070269 | TMEM260 | 5.053476 | 0.932347 | 0.004255 |
| ENSG00000104549 | SQLE | 7.208872 | 0.932566 | 0.002048 |
| ENSG00000092531 | SNAP23 | 6.063157 | 0.93365 | 0.002489 |
| ENSG00000075426 | FOSL2 | 3.575391 | 0.942459 | 0.040325 |
| ENSG00000184009 | ACTG1 | 5.683456 | 0.942787 | 0.002398 |
| ENSG00000150403 | TMCO3 | 5.926079 | 0.944941 | 0.002048 |
| ENSG00000035403 | VCL | 7.371297 | 0.945275 | 0.001725 |
| ENSG00000198624 | CCDC69 | 6.200573 | 0.945489 | 0.001907 |
| ENSG00000100558 | PLEK2 | 6.761641 | 0.946743 | 0.001594 |
| ENSG00000116161 | CACYBP | 5.931754 | 0.948167 | 0.002166 |
| ENSG00000071537 | SEL1L | 7.960664 | 0.949131 | 0.002 |
| ENSG00000110274 | CEP164 | 4.921205 | 0.952916 | 0.003601 |
| ENSG00000124942 | AHNAK | 6.252599 | 0.954514 | 0.001715 |
| ENSG00000165521 | EML5 | 3.894542 | 0.965495 | 0.019081 |
| ENSG00000067369 | TP53BP1 | 5.316814 | 0.970235 | 0.001904 |
| ENSG00000147416 | ATP6V1B2 | 8.254907 | 0.973902 | 0.001518 |
| ENSG00000140650 | PMM2 | 3.727409 | 0.990141 | 0.02516 |
| ENSG00000101104 | PABPC1L | 3.332738 | 0.995589 | 0.043404 |
| ENSG00000155850 | SLC26A2 | 5.45092 | 0.995873 | 0.0011 |
| ENSG00000102230 | PCYT1B | 6.237495 | 0.997008 | 0.00085 |
| ENSG00000044574 | HSPA5 | 8.282878 | 0.99833 | 0.001095 |
| ENSG00000169905 | TOR1AIP2 | 9.181014 | 0.999026 | 0.001307 |
| ENSG00000130513 | GDF15 | 8.85964 | 1.000826 | 0.001159 |
| ENSG00000151929 | BAG3 | 5.112944 | 1.001641 | 0.001448 |
| ENSG00000173281 | PPP1R3B | 6.536975 | 1.005601 | 0.000686 |
| ENSG00000166147 | FBN1 | 3.924932 | 1.006258 | 0.012957 |
| ENSG00000082641 | NFE2L1 | 6.827287 | 1.008723 | 0.000612 |
| ENSG00000102265 | TIMP1 | 3.644348 | 1.008891 | 0.031111 |
| ENSG00000100219 | XBP1 | 5.406715 | 1.01837 | 0.000826 |

| | | | | |
|---|---|---|---|---|
| ENSG00000151208 | DLG5 | 5.225762 | 1.020862 | 0.000902 |
| ENSG00000080031 | PTPRH | 4.52181 | 1.021103 | 0.002723 |
| ENSG00000270055 | | 3.619299 | 1.043086 | 0.017653 |
| ENSG00000121060 | TRIM25 | 4.879517 | 1.043364 | 0.001373 |
| ENSG00000152689 | RASGRP3 | 5.39937 | 1.043864 | 0.000511 |
| ENSG00000109814 | UGDH | 3.923867 | 1.052769 | 0.006831 |
| ENSG00000052802 | MSMO1 | 6.538679 | 1.059595 | 0.000328 |
| ENSG00000143515 | ATP8B2 | 5.06243 | 1.070078 | 0.00056 |
| ENSG00000166068 | SPRED1 | 3.174856 | 1.086953 | 0.036872 |
| ENSG00000198692 | EIF1AY | 6.350794 | 1.105137 | 0.000139 |
| ENSG00000135269 | TES | 4.397026 | 1.106944 | 0.001102 |
| ENSG00000138413 | IDH1 | 4.982712 | 1.107995 | 0.000359 |
| ENSG00000074842 | MYDGF | 3.572245 | 1.110727 | 0.014121 |
| ENSG00000120217 | CD274 | 4.1302 | 1.115891 | 0.002 |
| ENSG00000136603 | SKIL | 4.360075 | 1.118051 | 0.001274 |
| ENSG00000133318 | RTN3 | 7.751166 | 1.128463 | 0.000107 |
| ENSG00000164111 | ANXA5 | 4.643894 | 1.131706 | 0.00048 |
| ENSG00000134070 | IRAK2 | 4.087793 | 1.145141 | 0.001518 |
| ENSG00000080824 | HSP90AA1 | 10.54948 | 1.167182 | 0.000139 |
| ENSG00000100600 | LGMN | 3.988343 | 1.167502 | 0.001839 |
| ENSG00000136048 | DRAM1 | 5.291826 | 1.174654 | 6.6E-05 |
| ENSG00000172889 | EGFL7 | 2.932455 | 1.175449 | 0.046588 |
| ENSG00000105329 | TGFB1 | 4.208301 | 1.189161 | 0.000813 |
| ENSG00000187240 | DYNC2H1 | 3.807732 | 1.19324 | 0.002 |
| ENSG00000124491 | F13A1 | 3.061732 | 1.200058 | 0.026758 |
| ENSG00000124171 | PARD6B | 3.410701 | 1.203505 | 0.007225 |
| ENSG00000165271 | NOL6 | 5.52577 | 1.230613 | 2E-05 |
| ENSG00000078269 | SYNJ2 | 3.503638 | 1.235993 | 0.003746 |
| ENSG00000122862 | SRGN | 4.19112 | 1.25246 | 0.000347 |
| ENSG00000143878 | RHOB | 5.861825 | 1.259718 | 8.14E-06 |
| ENSG00000110172 | CHORDC1 | 6.400267 | 1.266058 | 6.72E-06 |
| ENSG00000276248 | | 2.614099 | 1.307039 | 0.032254 |
| ENSG00000099204 | ABLIM1 | 5.153161 | 1.307681 | 6.72E-06 |
| ENSG00000108854 | SMURF2 | 3.555777 | 1.311973 | 0.001448 |
| ENSG00000115977 | AAK1 | 7.544154 | 1.317763 | 2.11E-06 |
| ENSG00000197892 | KIF13B | 5.412193 | 1.322027 | 2.89E-06 |
| ENSG00000116016 | EPAS1 | 4.22565 | 1.325738 | 8.82E-05 |
| ENSG00000124762 | CDKN1A | 4.795758 | 1.329192 | 8.95E-06 |
| ENSG00000178381 | ZFAND2A | 4.285522 | 1.353051 | 4.74E-05 |
| ENSG00000004478 | FKBP4 | 6.626593 | 1.354538 | 1.06E-06 |
| ENSG00000163435 | ELF3 | 3.533806 | 1.357518 | 0.000841 |
| ENSG00000230453 | ANKRD18B | 4.687825 | 1.366825 | 7.99E-06 |
| ENSG00000204387 | SNHG32 | 5.269065 | 1.369789 | 1.67E-06 |
| ENSG00000157827 | FMNL2 | 3.404906 | 1.377295 | 0.001102 |
| ENSG00000071575 | TRIB2 | 3.467876 | 1.380105 | 0.001012 |
| ENSG00000138166 | DUSP5 | 3.131741 | 1.413339 | 0.002353 |

| | | | | |
|---|---|---|---|---|
| ENSG00000158079 | PTPDC1 | 2.583527 | 1.414843 | 0.016918 |
| ENSG00000135046 | ANXA1 | 2.215552 | 1.415565 | 0.039644 |
| ENSG00000120889 | TNFRSF10B | 4.632691 | 1.430273 | 2.81E-06 |
| ENSG00000138061 | CYP1B1 | 3.996962 | 1.44578 | 3.93E-05 |
| ENSG00000198380 | GFPT1 | 6.746248 | 1.446074 | 1.06E-07 |
| ENSG00000120694 | HSPH1 | 7.595225 | 1.449715 | 1.1E-07 |
| ENSG00000110848 | CD69 | 3.248478 | 1.451371 | 0.001012 |
| ENSG00000197586 | ENTPD6 | 4.254531 | 1.463377 | 9.71E-06 |
| ENSG00000250337 | PURPL | 2.033645 | 1.466187 | 0.036595 |
| ENSG00000264175 | MIR3189 | 2.690135 | 1.470943 | 0.007569 |
| ENSG00000178607 | ERN1 | 5.2623 | 1.478003 | 1.1E-07 |
| ENSG00000100994 | PYGB | 4.628863 | 1.486239 | 1.06E-06 |
| ENSG00000178695 | KCTD12 | 1.900417 | 1.506848 | 0.044282 |
| ENSG00000118263 | KLF7 | 1.977121 | 1.532672 | 0.030206 |
| ENSG00000115425 | PECR | 2.249274 | 1.55755 | 0.019882 |
| ENSG00000135919 | SERPINE2 | 4.710971 | 1.57311 | 1.06E-07 |
| ENSG00000226562 | CYP4F26P | 2.173416 | 1.575774 | 0.016918 |
| ENSG00000033327 | GAB2 | 1.914881 | 1.582777 | 0.027995 |
| ENSG00000132196 | HSD17B7 | 2.415202 | 1.655801 | 0.0075 |
| ENSG00000242574 | HLA-DMB | 0.8555 | 1.661178 | 0.04269 |
| ENSG00000262001 | DLGAP1-AS2 | 3.087231 | 1.683616 | 0.000224 |
| ENSG00000010327 | STAB1 | 3.144458 | 1.695328 | 0.000301 |
| ENSG00000038427 | VCAN | 0.30217 | 1.732725 | 0.045755 |
| ENSG00000186481 | ANKRD20A5P | 0.847373 | 1.743808 | 0.031114 |
| ENSG00000120738 | EGR1 | 8.329744 | 1.744736 | 4.95E-11 |
| ENSG00000166927 | MS4A7 | 1.328776 | 1.771228 | 0.030706 |
| ENSG00000132002 | DNAJB1 | 7.738378 | 1.838664 | 1.43E-12 |
| ENSG00000187608 | ISG15 | 1.410874 | 1.882034 | 0.014121 |
| ENSG00000204388 | HSPA1B | 7.607862 | 2.036774 | 1.21E-15 |
| ENSG00000143674 | MAP3K21 | 0.600192 | 2.116474 | 0.030804 |
| ENSG00000173110 | HSPA6 | 3.792871 | 2.255705 | 2.54E-11 |
| ENSG00000099251 | HSD17B7P2 | 0.084555 | 2.30862 | 0.007871 |
| ENSG00000240457 | RN7SL472P | 1.466598 | 2.312627 | 0.000902 |
| ENSG00000204389 | HSPA1A | 6.967433 | 2.430663 | 1.34E-22 |
| ENSG00000156414 | TDRD9 | 1.462002 | 2.467771 | 0.00048 |

**Supplemental figure 1**

A B

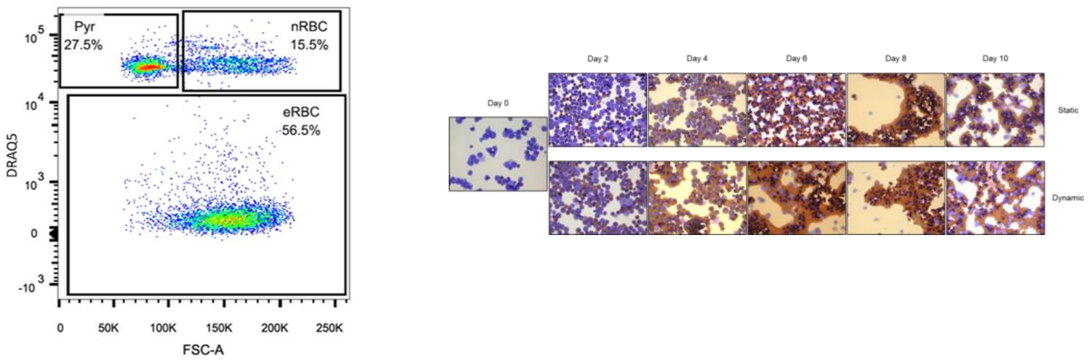


Supplemental figure 1. A) Gating strategy to determine cell enucleation. Cells were stained with DRAQ5 and plotted against forward scatter (FCS) as described in methods. The 3 detected populations are defined as pyrenocytes (PYR: $DRAQ5^{+}/FSC^{low}$), nucleated red blood cells (nRBC: $DRAQ5^{+}/FSC^{high}$) and enucleated red blood cells (eRBC: $DRAQ5^{-}/FSC^{high}$). B) Representative cytospins of erythroid precursors during 10 days of differentiation. Cells from static (top) and dynamic (bottom) conditions were cytospinned and stained with O-dianizidine to visualise heme (brown, hemoglobin) and water-soluble general cell dyes to visualise DNA content (purple) and cytoplasm (light blue). Imaged with 50×objective as described in material and methods).

**Supplemental figure 2**

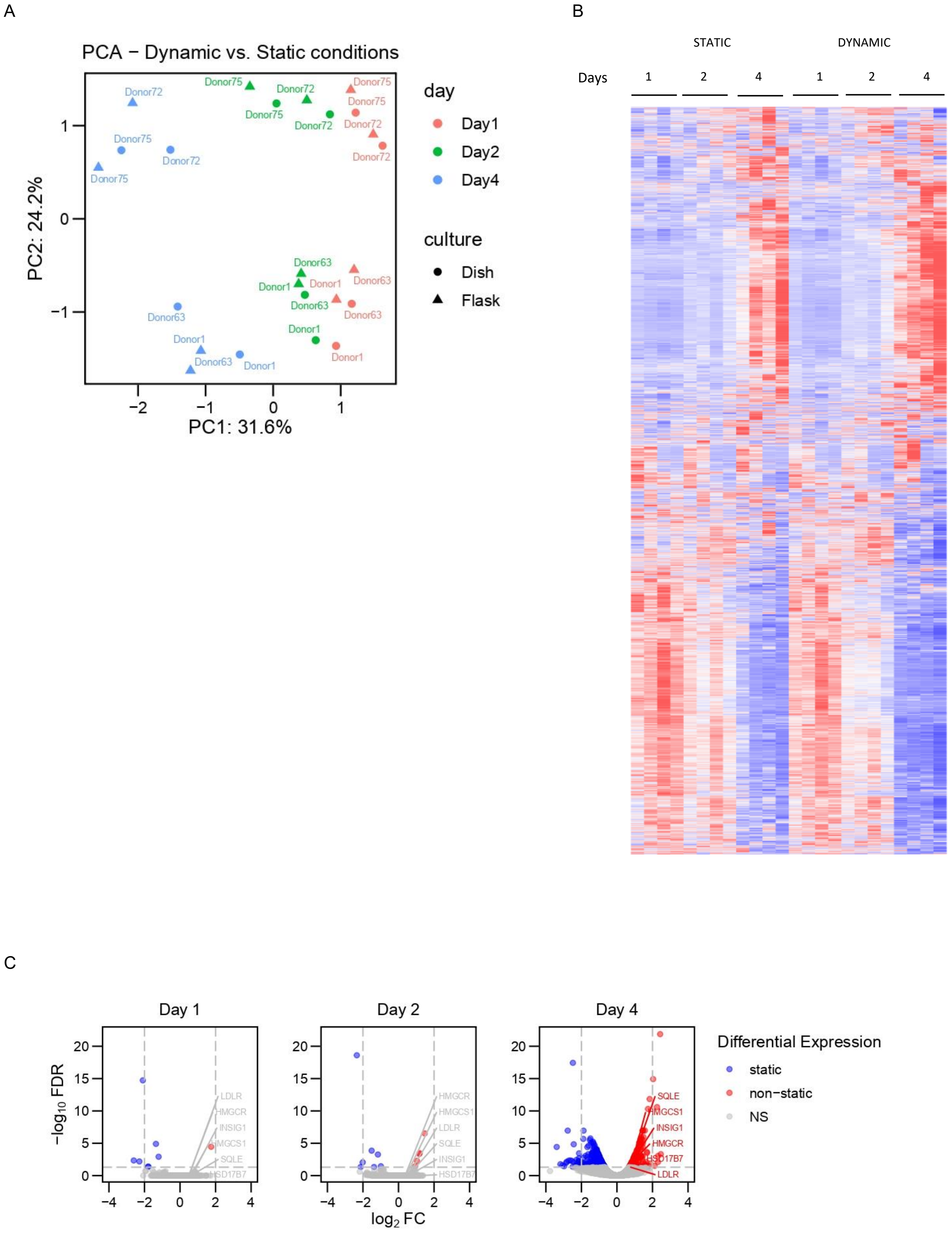

Supplemental figure 2. A) Principal component analysis derived from RNAseq of samples differentiated in static and dynamic conditions. X-axis inversely correlates to differentiation rate, y-axis represents variations between donors. B) Heatmap of transcripts differentially expressed during the first 4 days of differentiation in static (left) and dynamic (right) conditions. C) Volcano plots of transcripts differentially expressed during the first 4 days of differentiation in static (blue) and dynamic (red) conditions. Transcripts involved in the cholesterol metabolism are annotated.

### Supplemental figure 3

A

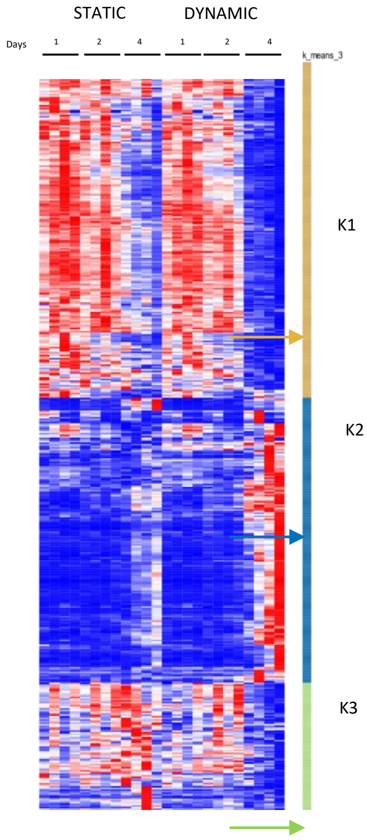


| Biological process | P-value | Adjusted P-value | Genes |
|---|---|---|---|
| DNA Unwinding Involved In DNA Replication (GO:0006268) | 0,0000000000002 | 0,00000000024 | RECQL4;GINS2;GINS3;GINS4;MCM3;MCM4;MCM5;MCM6;MCM2 |
| Double-Strand Break Repair Via Break-Induced Replication (GO:0000727) | 0,0000000000042 | 0,00000000239 | GINS2;GINS4;MCM3;MCM4;MCM5;MCM6;MCM2 |
| DNA-templated DNA Replication (GO:0006261) | 0,0000000000072 | 0,00000000274 | POLD3;RECQL4;PRIM2;GINS2;POLD1;GINS3;GINS4;MCM3;MCM4;MCM5;MCM6;MCM2 |
| DNA Duplex Unwinding (GO:0032508) | 0,0000000003568 | 0,00000010170 | RECQL4;GINS2;GINS3;GINS4;MCM3;MCM4;MCM5;MCM6;MCM2 |
| DNA Replication (GO:0006260) | 0,0000000048052 | 0,00000109558 | POLD3;RECQL4;DUT;POLD1;CDK2;PCLAF;MCM4;MCM5;MCM6;MCM2 |
| Regulation Of DNA-templated DNA Replication (GO:0090329) | 0,0000000144234 | 0,00000274044 | CDT1;MCM3;MCM4;MCM5;MCM6;MCM2 |
| Double-Strand Break Repair Via Homologous Recombination (GO:0000724) | 0,0000000261293 | 0,00000425534 | RECQL4;GINS2;XRCC2;XRCC3;GINS4;TONSL;MCM3;MCM4;MCM5;MCM6;MCM2 |
| Mitotic DNA Replication (GO:1902969) | 0,0000000300865 | 0,00000428732 | GINS3;MCM3;MCM4;MCM6;MCM2 |
| DNA Metabolic Process (GO:0006259) | 0,0000000750015 | 0,00000950019 | PRIM2;DUT;PARP1;XRCC2;XRCC3;RECQL4;MSH6;POLD3;POLD1;CDK2;PCLAF;MCM3;MCM4;MCM5;MCM6;MCM2 |
| Regulation Of DNA-templated DNA Replication Initiation (GO:0030174) | 0,0000000929442 | 0,00001059564 | MCM3;MCM4;MCM5;MCM6;MCM2 |

| Biological process | P-value | Adjusted P-value | Genes |
|---|---|---|---|
| Sterol Biosynthetic Process (GO:0016126) | 0.000000003 | 0.000001601 | SQLE;HMGCS1;INSIG1;HMGCR;HSD17B7 |
| Secondary Alcohol Biosynthetic Process (GO:1902653) | 0.000000157 | 0.000037106 | HMGCS1;INSIG1;HMGCR;HSD17B7 |
| Cholesterol Biosynthetic Process (GO:0006695) | 0.000000186 | 0.000037106 | HMGCS1;INSIG1;HMGCR;HSD17B7 |
| Cholesterol Metabolic Process (GO:0008203) | 0.000000247 | 0.000037106 | SQLE;HMGCS1;INSIG1;HMGCR;HSD17B7 |
| Regulation Of Protein Secretion (GO:0050708) | 0.000055852 | 0.006702296 | TMED10;TGFB1;STXBP5;HMGCR |
| Response To Sterol (GO:0036314) | 0.000182593 | 0.018259317 | TGFB1;INSIG1 |
| Negative Regulation Of Nitrogen Compound Metabolic Process (GO:0051172) | 0.000222880 | 0.019103994 | TGFB1;LDLR |
| Membrane Protein Intracellular Domain Proteolysis (GO:0031293) | 0.000315265 | 0.023644853 | TGFB1;HM13 |
| Positive Regulation Of Isotype Switching (GO:0045830) | 0.000423295 | 0.025475935 | TGFB1;TP53BP1 |
| Positive Regulation Of Secretion By Cell (GO:1903532) | 0.000450152 | 0.025475935 | TMED10;TGFB1;STXBP5 |

| Biological process | P-value | Adjusted P-value | Genes |
|---|---|---|---|
| Innate Immune Response In Mucosa (GO:0002227) | 0,000004 | 0,002127 | H2BC12;H2BC8;H2BC21;H2BC11 |
| Nucleosome Assembly (GO:0006334) | 0,000009 | 0,002127 | H2BC14;H2BC8;H2BC21;H2BC3;H2BC11 |
| Mucosal Immune Response (GO:0002385) | 0,000012 | 0,002127 | H2BC12;H2BC8;H2BC21;H2BC11 |
| protein-DNA Complex Assembly (GO:0065004) | 0,000013 | 0,002127 | H2BC14;H2BC8;H2BC21;H2BC3;H2BC11 |
| Nucleosome Organization (GO:0034728) | 0,000043 | 0,005495 | H2BC14;H2BC8;H2BC21;H2BC3;H2BC11 |
| Antibacterial Humoral Response (GO:0019731) | 0,000118 | 0,012644 | H2BC12;H2BC8;H2BC21;H2BC11 |
| Antimicrobial Humoral Immune Response Mediated By Antimicrobial Peptide (GO:0061844) | 0,000353 | 0,032516 | H2BC12;H2BC8;H2BC21;H2BC11 |
| Defense Response To Bacterium (GO:0042742) | 0,000658 | 0,053071 | H2BC12;H2BC8;H2BC21;STAB1;H2BC11;AQP1 |
| Defense Response To Gram-positive Bacterium (GO:0050830) | 0,000976 | 0,067463 | H2BC12;H2BC8;H2BC21;H2BC11 |
| Odontogenesis (GO:0042476) | 0,001059 | 0,067463 | OSR2;ROGDI;AQP1 |

B

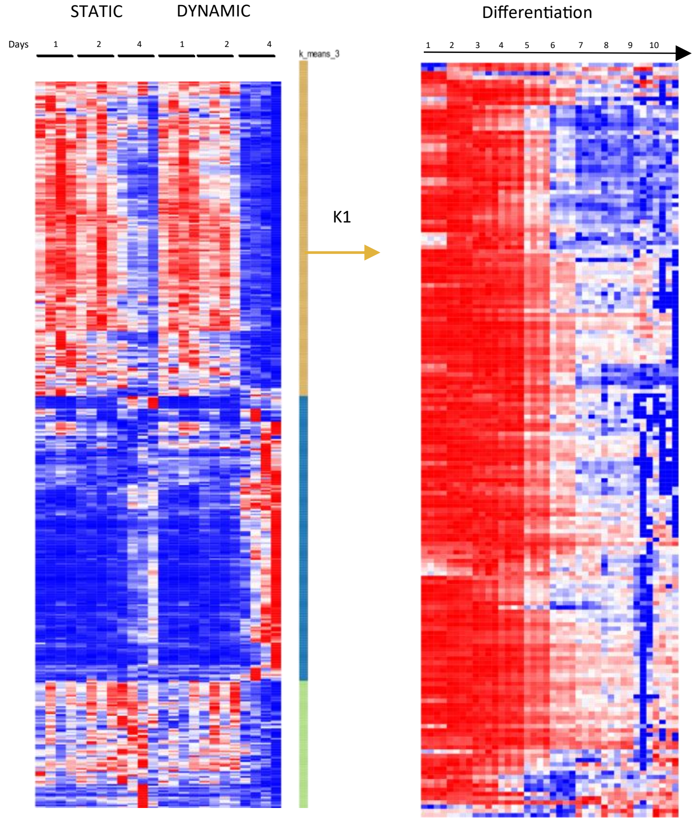


(Heshusius et al, 2019)

C

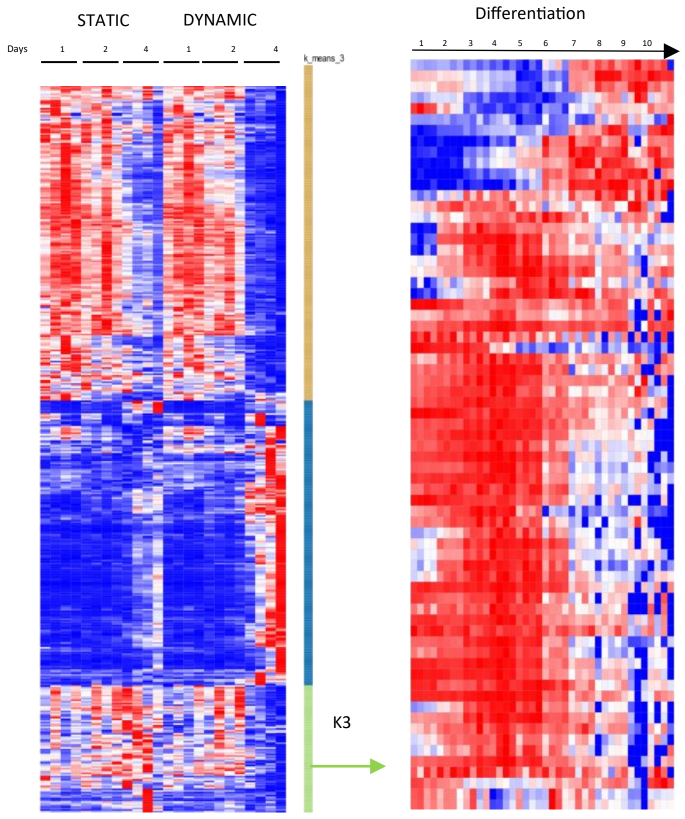


(Heshusius et al, 2019)

Supplemental figure 3. A) Gene ontology analysis (EnRicHR) of genes within Kmeans cluster1 (K1) and Kmeans cluster 3 (K3) from figure 3A. The first 10 biological processes have been selected according to the adjusted p-value. B and C) The gene identifiers of the upregulated RNAs of k-means cluster 1 and 3 from figure 3A were extracted and their expression dynamics through complete differentiation of EBL to enucleated reticulocytes as previously published by Heshusius[5] was datamined and determined (n=4 different donors, time points as indicated).

**Supplemental figure 4**

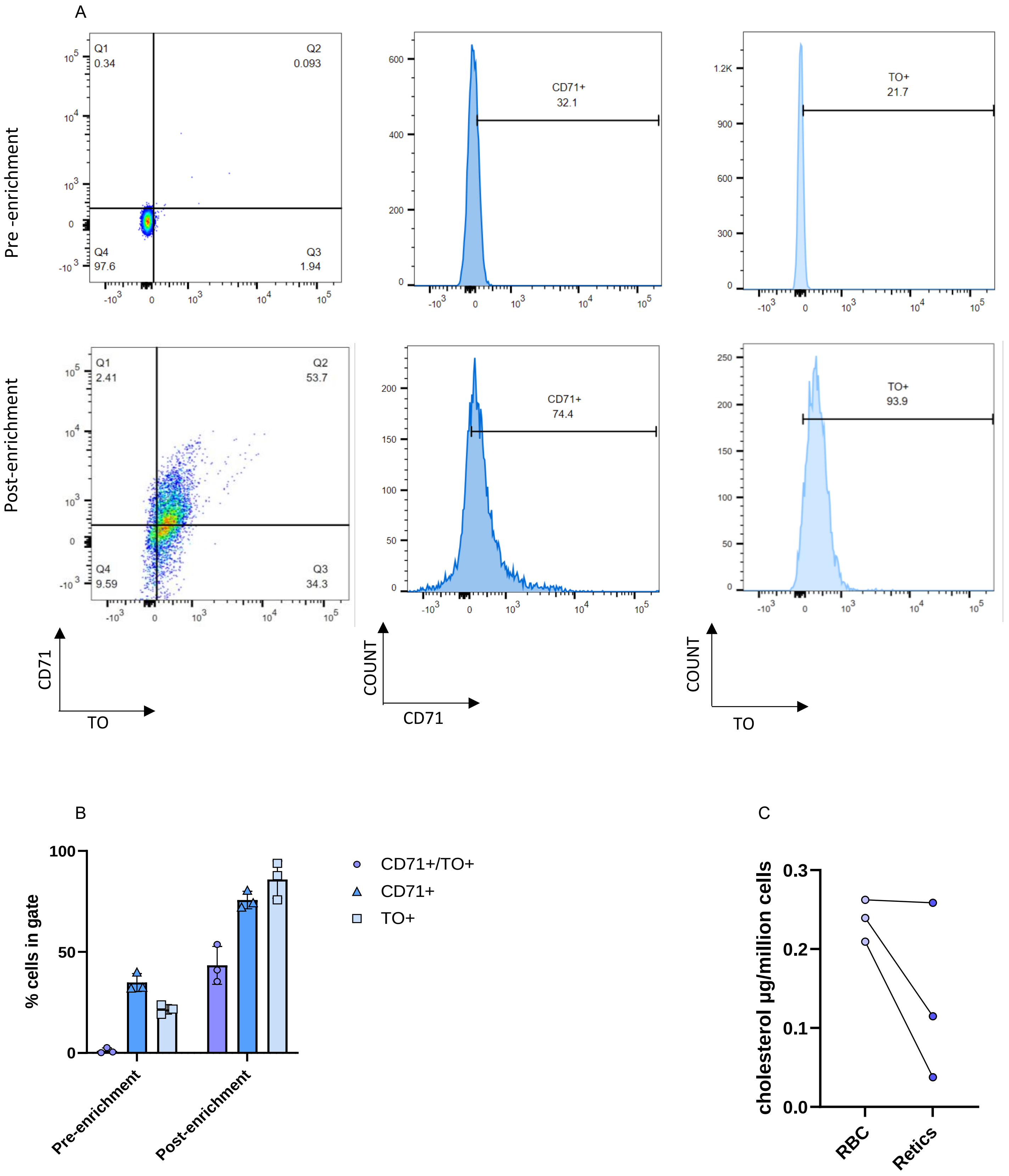

Supplemental figure 4. Evaluation of the immunomagnetic (MACS) reticulocyte enrichment A) Representative example of gating strategy for RBC (pre enrichment) and reticulocytes (post enrichment). RBC and reticulocytes were stained with TO and anti-CD71 and analysed by flow cytometry.
B) TO and CD71 expression was averaged for 3 donors (n=3) by quantifying the percentage of cells within the respective gates, for RBC (pre enrichment) and reticulocytes (post enrichment). C) RBC vs reticulocytes cholesterol concentration (μg cholesterol/million cells).

**Supplemental figure 5**

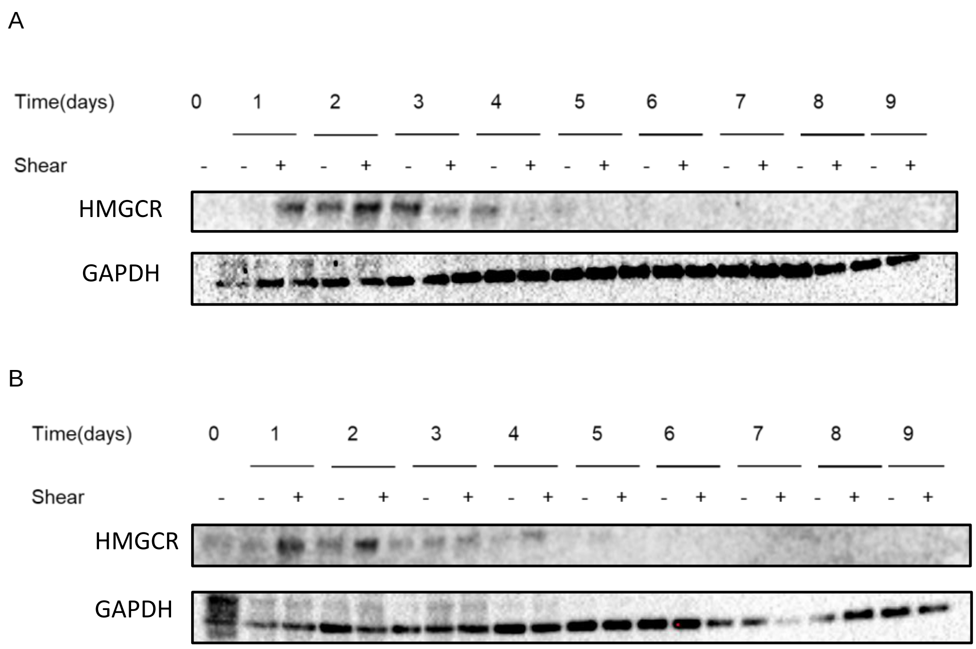


Supplemental figure 5. A and B) Representative examples of HMGCR western blot for two different donors during differentiation in static and dynamic conditions. GAPDH staining was used as loading control. Data of figure 5B derived from the normalised (according to GAPDH) and averaged values of the 3 blots showed (Figure 5A and supplemental figure 5).

**Supplemental figure 6**

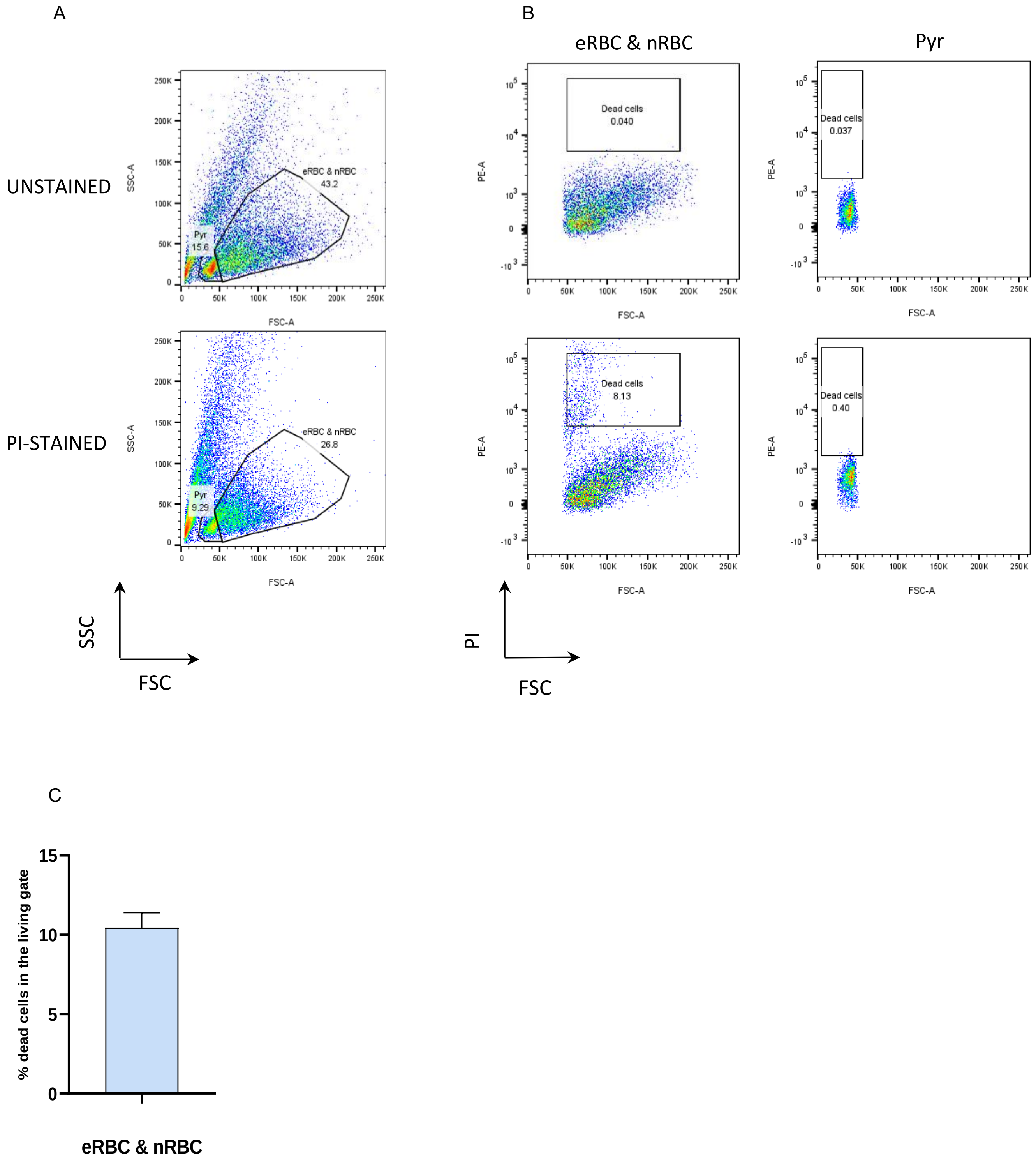


Supplemental figure 6. A) Representative example of propidium iodide (PI) gating strategy used to confirm viability of cells detected in living gate (Figure 6). Forward scatter (FSC) vs side scatter (SSC) were plotted and cells gated as follows: Pyr (pyrenocytes): $FSC^{low}/SSC^{low}$; eRBC & nRBC (enucleated RBC: reticulocytes and nucleated RBC: EBL): $FSC^{high}/SSC^{high}$. B) PI vs FSC were plotted to determine the % of living cells in gates. A and B) Unstained samples (top); PI-stained samples (bottom). C) Percentage of dead eRBC & nRBC in living gate averaged for 5 donors.
